# Large-Scale Genome-Wide Optimization and Prediction of the Cre Recombinase System for Precise Genome Manipulation in Mice

**DOI:** 10.1101/2024.06.14.599022

**Authors:** Valerie Erhardt, Elli Hartig, Kristian Lorenzo, Hannah R. Megathlin, Basile Tarchini, Vishnu Hosur

## Abstract

The Cre-Lox recombination system is a powerful tool in mouse genetics, offering spatial-temporal control over gene expression and facilitating the large-scale generation of conditional knockout mice. Its versatility also extends to other research models, such as rats, pigs, and zebrafish. However, the Cre-Lox technology presents a set of challenges that includes high costs, a time-intensive process, and the occurrence of unpredictable recombination events, which can lead to unexpected phenotypic outcomes. To better understand factors affecting recombination, we embarked on a systematic and genome-wide analysis of Cre-mediated recombination in mice. To ensure uniformity and reproducibility, we generated 11 novel strains with conditional alleles at the *ROSA26* locus, utilizing a single inbred mouse strain background, C57BL/6J. We examined several factors influencing Cre-recombination, including the inter-*loxP* distance, mutant *loxP* sites, the zygosity of the conditional alleles, chromosomal location, and the age of the breeders. We discovered that the selection of the Cre-driver strain profoundly impacts recombination efficiency. We also found that successful and complete recombination is best achieved when *loxP* sites are spaced between 1 to 4 kb apart, with mutant *loxP* sites facilitating recombination at distances of 1 to 3 kb. Furthermore, we demonstrate that complete recombination does not occur at an inter-*loxP* distance of ≥ 15 kb with wildtype *loxP* sites, nor at a distance of ≥ 7 kb with mutant *lox71/66* sites. Interestingly, the age of the Cre-driver mouse at the time of breeding emerged as a critical factor in recombination efficiency, with best results observed between 8 and 20 weeks old. Moreover, crossing heterozygous floxed alleles with the Cre-driver strain resulted in more efficient recombination than using homozygous floxed alleles. Lastly, maintaining an inter-*loxP* distance of 4 kb or less ensures efficient recombination of the conditional allele, regardless of the chromosomal location. While CRISPR/Cas has revolutionized genome editing in mice, Cre-Lox technology remains a cornerstone for the generation of sophisticated alleles and for precise control of gene expression in mice. The knowledge gained here will enable investigators to select a Cre-Lox approach that is most efficient for their desired outcome in the generation of both germline and non-germline mouse models of human disease, thereby reducing time and cost of Cre-Lox technology-mediated genome modification.

## Introduction

Cre-Lox recombination systems are versatile mouse genetics tools, permitting spatial-temporal control of gene expression^1–6^. Since their development in the 1980s^7,8^, they have become a widely accepted and extensively used method for genome editing, paving the way for new opportunities across various research fields. The applications of this increasingly potent technology are diverse, encompassing the generation of targeted deletions, translocations, or inversions, including double-floxed inverse orientation^9^. Cre-Lox applications also extend to trans-chromosomal recombination for generating deletions or duplications^10^, the Brainbow system^11,12^, which allows individual neurons to be marked by different colors, and to a growing array of inducible systems^13,14^. The advent of Cre-Lox technology has expedited the production of conditional knockout mice on a large scale^15,16^. Its utility is not limited to mice but extends to other research model animals such as rats, pigs, and zebrafish. The enduring relevance, interest, and continuing advancements of the Cre-Lox system are underscored by the high volume of publications using it. Since the beginning of 2024, 376 preprints related to ‘Cre-Lox technology’ have been shared on BioRxiv between January 1^st^ and June 1st, 2024. This underscores the sustained interest and continuous advancements in the field.

In its most basic form, the Cre-Lox system in mice consists of a Cre driver, which provides the Cre protein, a Cre reporter (e.g., lacZ or a fluorescent reporter) and a floxed allele^8^. The floxed allele contains a target sequence—the region of Cre-mediated recombination—enclosed by a pair of directional *loxP* sites, each approximately 34 bp in length. The Cre protein binds the pair of *loxP* sites and initiates recombination between them^17,18^. The outcome of the recombination depends on the orientation of the *loxP* sites; recombination between *loxP* sites oriented in the opposite direction results in an inversion of the intervening sequence, while recombination between *loxP* sites oriented in the same direction results in deletion of the intervening sequence. Despite the proven value of the Cre-Lox technology, it comes with a hefty price tag and is time-consuming—the cost of generating conditional alleles is approximately $16,000 to $18,000, and the process takes around 7-9 months (Sr. Study Director, Genetic Engineering Technologies, The Jackson Laboratory, personal communication). Moreover, it is associated with specific biological challenges^19–24^, including mosaicism of recombination within target tissues, which can lead to unexpected phenotypic outcomes (Fig. 1a). Even so, genetic mosaicism, a condition in which a multicellular organism possesses two or more populations of cells with different genetic makeup, is not necessarily an undesirable outcome for all applications. Mosaicism occurs in several human pathologies, including cancer, and therefore being able to predictably generate mosaic animals for translational research is as important as generating animals with a uniform genotype in every cell.

**Figure 1.**
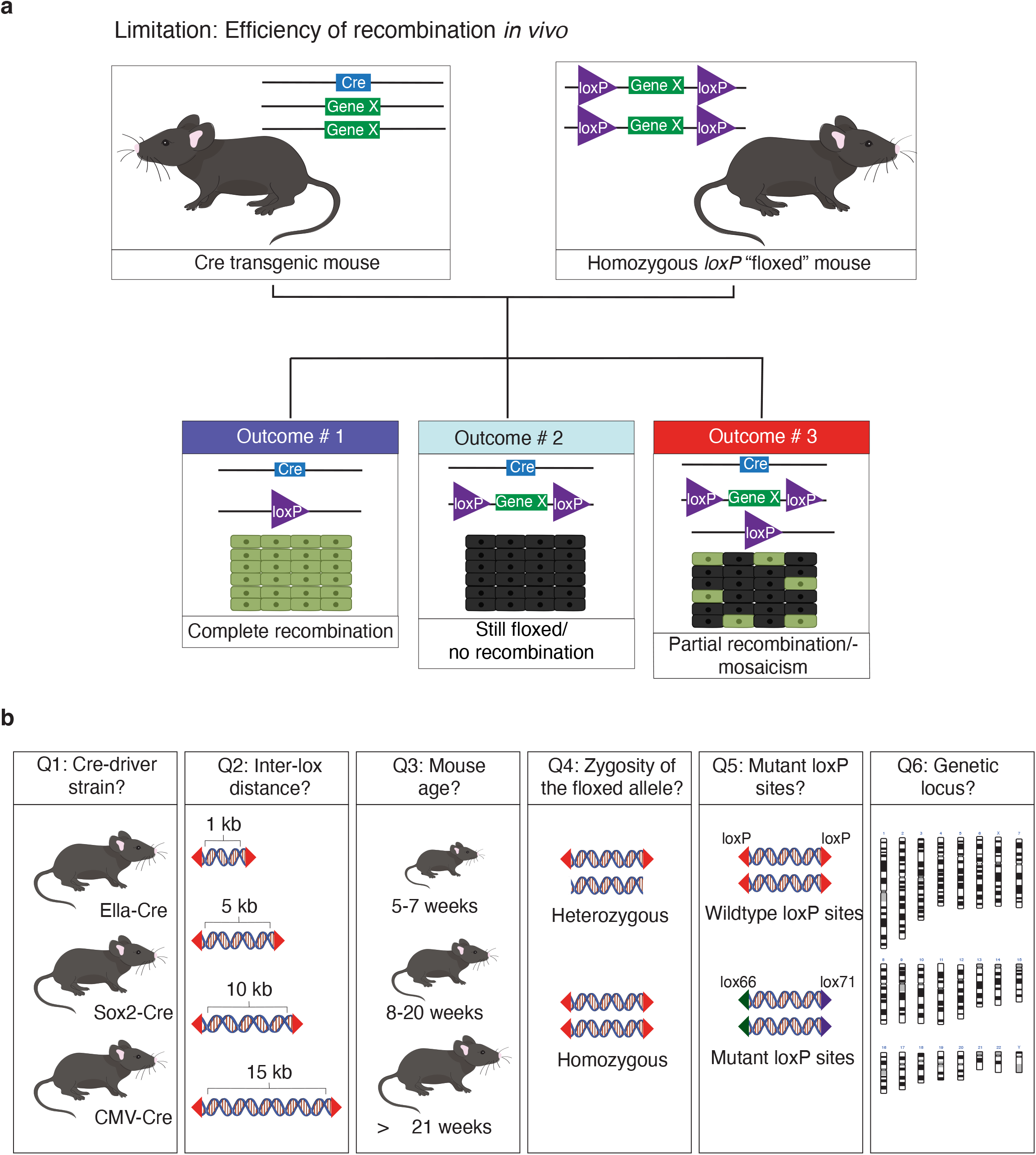
Optimizing Cre-Lox recombination: factors affecting efficiency and reliability. (a) The Cre-Lox system, consisting of the enzyme Cre and its specific DNA sequence lox, enables spatial-temporal control over gene expression in mice. It is widely used in research, allowing researchers to study the function of specific genes with precise control over expression time and location, deepening our understanding of biological processes. Cre-Lox technology has a notable limitation—the unpredictability of efficiency of Cre-mediated recombination and mosaicism (i.e., incomplete recombination—the targeted genetic changes do not occur uniformly across the targeted tissue). Although a high recombination efficiency is in most cases the desired outcome, for some biological questions, genetic mosaicism is of physiological relevance. Our lack of understanding of how various considerations affect recombination efficiency, especially when using tissue-specific Cre drivers, undermines the reliability of the Cre-Lox system, leading to variable experimental outcomes that may not align optimally with the investigator’s intention. (b) We systematically examined the role of various factors, including the Cre-driver strain, inter-*loxP* distance, Cre breeder age at the time of mating, zygosity of the floxed allele (heterozygous versus homozygous floxed allele), wildtype versus mutant *loxP* sites, and the various genetic loci of floxed alleles across the genome on Cre-recombination efficiency and mosaicism.

Emerging literature suggests that the efficiency of Cre-recombination can be influenced by the Cre-driver strain^23,25^ as well as the inter-*loxP* distance; in mouse embryonic stem cell lines, Cre-Lox recombination has been shown to be effective for floxed regions of up to a few centimorgans (cM), and this efficiency has been shown to decrease over increasing genetic distances (2 cM to 60 cM)^26,27^. However, the genetic distances for effective recombination *in vivo* in mice are poorly understood^11,28–30^. Furthermore, several recent studies highlight variable, often incomplete, and mosaic recombination with inter-*loxP* distances in the kilobase range^31–35^.

Genomic context, specifically the genetic locus targeted for recombination, is another significant factor. Several studies, including recent investigations using multiple Cre drivers^3,31,35–38^, have emphasized this point. For instance, using Cre mice (*R26cre*-*ER^T^*), Vooijs et al. showed significant differences in recombination frequency among the four loci examined in the study—*Rb*, *ROSA26*, *Brca2,* and *p53*. Similarly, Long et al. observed significant differences in recombination frequencies following the crossing to either LysM -cre or pCX-NLS-cre mice of three strains with three reporters (Z/AP, Z/EG, and R26R-EYFP) located in different chromosomal regions. In addition, Luo et al. found that following mating of female *Emx1*-cre; *Wwp1^flox/flox^; Wwp2 ^flox/flox^* mice with male *Wwp1 ^flox/flox^*; *Wwp2 ^flox/flox^* mice, recombination was observed at the *Wwp2* locus, but not at the *Wwp1* locus.

Despite the strong evidence linking recombination efficiency to various factors—the type of *loxP* sites (wild-type or mutant), the zygosity of the floxed allele (which can be either homozygous or heterozygous), and the age of the Cre-breeder strain—no study has yet systematically investigated the impact of these factors at a genome-wide scale or at a fixed locus in mice. Thus, we undertook an extensive genome-wide analysis of Cre-mediated recombination at 12 different loci across different chromosomes. A major challenge, specifically with respect to investigating the inter-*loxP* distance, has been the lack of technology to precisely integrate *loxP*-flanked transgenes in the range of 1 kb to 15 kb; currently available CRISPR/Cas9 technology^16,39^ is slow and difficult, because CRISPR/Cas9-mediated homology-directed repair is variable and inefficient for insertion of transgenes greater than 1-3 kb. Furthermore, insertion of *loxP* sites is also challenging with CRISPR/Cas9^16^. We have therefore used a novel high-efficiency *Bxb1* recombinase system^40–42^ to integrate, separately, 11 constructs with different inter-*loxP* site lengths into the *ROSA26* locus to uniformly, efficiently, and rapidly generate 11 novel floxed strains. The strains generated, the inter-*loxP* distance, and the type of *loxP* sites are summarized in Table 1. Importantly, the generation of these floxed alleles at a fixed locus has allowed us to determine the impact of inter-*loxP* distance, the Cre-driver strain and age at time of breeding, the zygosity of the floxed allele, and the type of *loxP* sites (wildtype vs. mutant) on Cre-recombination efficiency.

**Table 1.** List of the various floxed strains generated at a fixed locus—*ROSA26*—with either wildtype or mutant *loxP* sites. Two more strains were generated, each with a different inter-*loxP* distance at the same locus— *Rhbdf1*.

| Strain name | Inter-lox distance | Type of *loxP* |
| --- | --- | --- |
| *Rosa*-0.8 kb | 0.8 kb | Wildtype *loxP* sites |
| *Rosa*-2.25 kb | 2.25 kb | Wildtype *loxP* sites |
| * *Rosa* -4 kb | 4 kb | Wildtype *loxP* sites |
| * *Rosa* -8 kb | 8 kb | Wildtype *loxP* sites |
| * *Rosa* -10 kb | 10 kb | Wildtype *loxP* sites |
| * *Rosa* -12 kb | 12 kb | Wildtype *loxP* sites |
| * *Rosa* -15 kb | 15 kb | Wildtype *loxP* sites |
| * *Rosa-DIO* -1.1 kb | 1.1 kb | Double-Floxed Inverted |
| * *Rosa-DIO* -2.2 kb | 2.2 kb | Double-Floxed Inverted |
| * *Rosa-DIO* -5.3 kb | 5.3 kb | Double-Floxed Inverted |
| * *Rosa-DIO* -6.9 kb | 6.9 kb | Double-Floxed Inverted |
| * *Rosa-DIO* -8.1 kb | 8.1 kb | Double-Floxed Inverted |
| *Rhbdf1*-2.9 kb | 2.9 kb | *Lox71/Lox66* |
| *Rhbdf1*-6.9 kb | 6.9 kb | *Lox71/Lox66* |

In this study, we conducted a comprehensive analysis (Fig. 1b), focusing on Cre-mediated recombination and mosaicism in mice, examining a variety of factors that could potentially influence its efficiency. Our findings can be summarized as follows: a) The choice of Cre-driver strain plays a pivotal role in recombination efficiency, irrespective of the inter-*loxP* distance, b) recombination is most successful when *loxP* sites are separated by distances ranging from 1 to 4 kb, or c) 1 to 3 kb when working with mutant *loxP* sites, d) the age of the Cre-driver mouse at the time of breeding also matters, with the most efficient recombination observed when the breeder is between 8 and 20 weeks old, e) crossing the heterozygous floxed allele with the Cre-driver mouse resulted in more efficient recombination than when using the homozygous floxed allele, f) wildtype *loxP* sites proved to be more efficient than mutant *loxP* sites, and g) lastly, maintaining an inter-*loxP* (wildtype) distance of 4 kb or less ensures efficient recombination of the conditional allele, regardless of the genomic locus. By addressing various factors, we have laid the foundation for more efficient generation of precise mouse models for a wide range of human diseases.

## Results

### Cre-driver strain and inter-*loxP* distance are key determinants of recombination and mosaicism

Here, we primarily explored the influence of Cre-driver strain on Cre-mediated recombination and mosaicism at a fixed locus—*ROSA26*. We evaluated three widely utilized Cre-driver strains with robust promoters for global gene knockout: 1) Ella-cre, 2) CMV-cre, and 3) Sox2-cre (Fig. 2a). The Ella-cre driver strain served as a positive control, based on published data indicating that this strain mediates widespread mosaicism^43^. Our analysis encompassed the distinct inter-*loxP* distances: 0.8 kb, 4 kb, 8 kb, 10 kb, 12 kb, and 15 kb. The workflow involves breeding female Cre driver mice with male *Rosa*-floxed mice to produce F1 offspring. This breeding scheme was selected based on published data indicating that the Cre transgene is active in the female germline^44^, and therefore, regardless of the inheritance of the transgene, all progeny from female Cre mice show evidence of Cre enzyme activity. Between 1 to 8 litters, consisting of a total of 8 to 55 offspring, were then genotyped to assess the percent of three possible outcomes: complete recombination, mosaicism, or no recombination (Fig. 2b).

**Figure 2.**
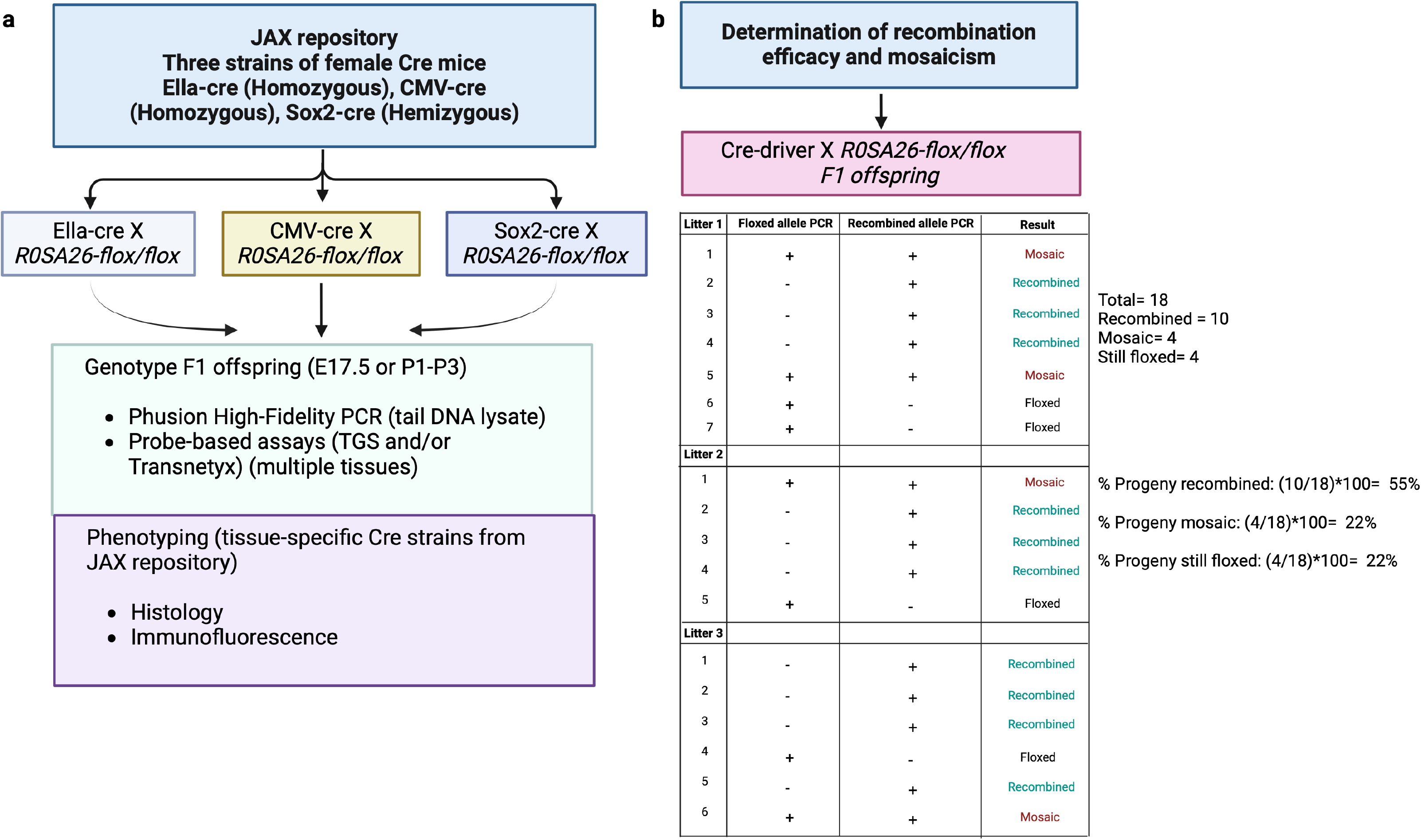
Overview of the pipeline and the methods used to assess the efficiency of recombination and mosaicism. (a) In this study, we implemented a thorough characterization pipeline comprising several crucial steps. Our initial focus was on characterizing three unique global Cre-driver strains, achieved by mating female Cre-driver strains with male *Rosa*-floxed strains. The resulting F1 offspring were genotyped using either standard PCR or probe-based assays. The genetic material for these assays was derived from tail tips of E17.5 stage embryos or postnatal P1 to P3 pups. For tissue-specific analysis of recombination and mosaicism, we conducted histological and immunohistochemical analyses using whole embryos from E15.5 to E17.5 or P2 pups, obtained from the mating of female floxed and male tissue-specific Cre-strains. (b)The percentage of progeny recombination was determined by dividing the number of litters showing complete recombination by the total number of litters and multiplying it by 100. The percentages of progeny mosaicism and floxed patterns were determined in a similar fashion.

For the inter-*loxP* distance of 0.8 kb (Fig. 3a), the Ella-cre x *Rosa*-0.8 kb flox combination resulted in 54% of the offspring exhibiting recombination, while 33% displayed mosaic traits. The CMV-cre x *Rosa*-0.8 kb flox combination had a slightly higher recombination rate of 57%, with 43% showing mosaic patterns. Interestingly, 13% of the offspring in the Ella-cre x *Rosa*-0.8 kb flox combination retained the floxed state, a phenomenon absent in the CMV-cre x *Rosa*-0.8 kb flox combination. The Sox2-cre x *Rosa*-0.8 kb flox combination, however, demonstrated a higher recombination rate of 79%, with only 16% showing mosaic patterns and a mere 5% remaining floxed (Fig. 3b and Supplementary Fig. 1a). Further analysis was conducted to confirm that the mosaicism observed in tail DNA lysate from mosaic offspring was representative of the entire mouse. We employed probe-based assays to examine the following tissues: liver, spleen, kidney, skin, and brain. The patterns of recombination, or mosaicism, were compared between the tail DNA lysate and the tissue-specific lysate. In most instances, we found a remarkable consistency in the patterns observed (Supplementary Fig. 1b). This consistency strongly suggests that tail DNA lysate could serve as a reliable predictor of the recombination/mosaic patterns in the entire organism.

**Figure 3.**
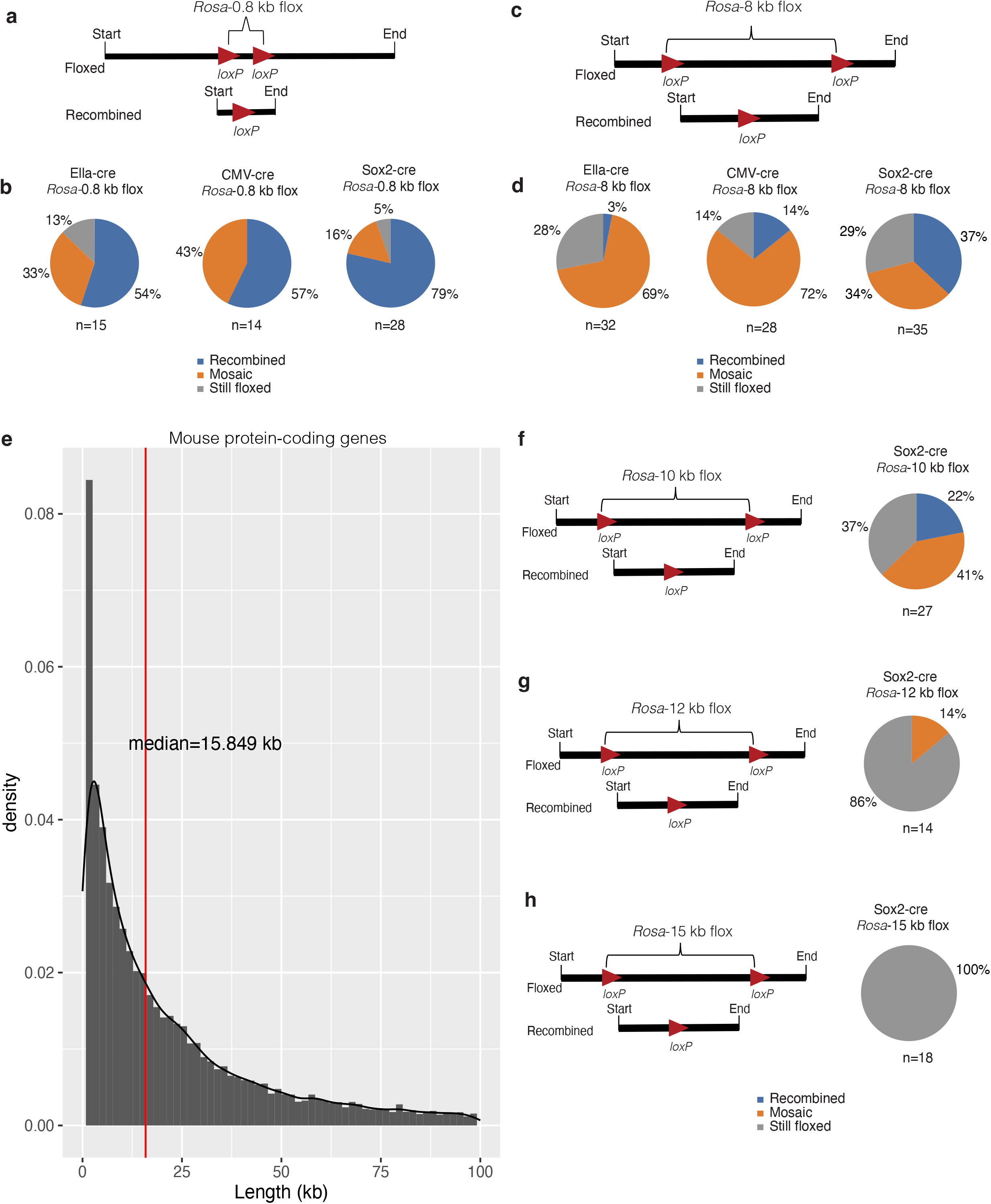
Cre-driver strain impact on recombination and mosaicism at different inter-*loxP* distances. (a) Schematic representation of a 0.8-kb floxed allele and a post-cre recombined allele. (b) Male homozygous floxed mice were bred with female Ella-cre, CMV-cre, or Sox2-cre mice. Note that Ella-cre and CMV-cre are homozygous, and Sox2-cre is hemizygous for the Cre allele. The F1 offspring were then genotyped using PCR from tail DNA to identify floxed and recombined alleles. Offspring with complete recombination were categorized as recombined, those with no recombination as still floxed, and those with both recombined and floxed alleles as mosaic. ‘n’ denotes the number of offspring genotyped. (c) Schematic representation of an 8-kb floxed allele and a post-cre recombined allele. Note that there is a 10-fold difference in the inter-*loxP* distance compared to the floxed allele analyzed in (a). (d) Homozygous floxed male mice were mated with either female Ella-cre, CMV-cre, or Sox2-cre mice, and the F1 offspring were genotyped to screen for floxed and recombined alleles. (e) The lengths of all protein-coding genes in mice are arranged in ascending order. Gene lengths varied widely from 0.066 kb to 2270 kb. The median length of protein-coding genes in mice is 15.8 kb. (f) Homozygous floxed male mice with a 10 kb inter-*loxP* distance were mated with Sox2-cre mice, and the F1 offspring were genotyped to screen for floxed and recombined alleles. Only 22% of the offspring showed complete recombination, while 41% were mosaic, and the remaining 37% showed no recombination. (g) Homozygous floxed male mice with a 12 kb inter-*loxP* distance were mated with Sox2-cre mice, and the F1 offspring were genotyped. None of the offspring showed complete recombination; however, 14% of the offspring were mosaic. (h) Homozygous floxed male mice with a 15 kb inter-*loxP* distance were mated with Sox2-cre mice, and the F1 offspring were genotyped using PCR from tail DNA to screen for floxed and recombined alleles. All the offspring had intact floxed alleles and showed no signs of recombination or mosaicism.

Next, we examined the impact of Cre-driver strain on recombination and mosaicism at an inter-*loxP* distance of ∼4 kb. The Ella-cre x *Rosa*-4 kb flox combination resulted in 16% of the offspring exhibiting recombination, while 60% displayed mosaic traits. The CMV-cre x *Rosa*-4 kb flox combination had a slightly higher recombination rate of 34%, with 53% showing mosaic patterns. Notably, 24% of the offspring in the Ella-cre x *Rosa*-4 kb flox combination retained the floxed state, compared to 13% in the CMV-cre x *Rosa*-4 kb flox combination. The Sox2-cre x *Rosa*-4 kb flox combination demonstrated a higher recombination rate of 73%, with only 27% showing mosaic patterns and 0% still floxed (Supplementary Fig. 2a).

We then investigated the impact of Cre-driver strain on Cre-mediated recombination and mosaicism at an inter-*loxP* distance of ∼8 kb (Fig. 3c). The Ella-cre x *Rosa*-8 kb flox combination resulted in a mere 3% of the offspring exhibiting recombination, while 69% displayed mosaic traits. The CMV-cre x *Rosa*-8 kb flox combination had a slightly higher recombination rate of 14%, with 72% showing mosaic patterns. Interestingly, 28% of the offspring in the Ella-cre x *Rosa*-8 kb flox combination retained the floxed state, compared to only 14% in the CMV-cre x *Rosa*-8 kb flox combination. The Sox2-cre x *Rosa*-8 kb flox combination demonstrated a higher recombination rate of 37%, with only 34% showing mosaic patterns and 29% remaining floxed (Fig. 3d).

Building upon these observations, our exploration delved further into understanding the nuances of genetic influence. Particularly intriguing were the findings when examining the tumor suppressor *Trp53* locus (Supplementary Fig. 2b). When we crossed Ella-cre with *Trp53*-8 kb flox, we observed that 19% of the offspring exhibited recombination. A higher recombination rate of 72% was noted when the CMV-cre strain was crossed with the *Trp53*-8 kb flox. However, the offspring of Sox2-cre and *Trp53*-8 kb flox demonstrated a higher recombination rate of 82%. In comparison to the *Rosa*-8 kb flox F1 offspring (Fig. 3d), the offspring of *Trp53*-8 kb flox showed higher recombination rates with all three Cre drivers (Supplementary Fig. 2b). This suggests that genetic loci can also impact Cre-mediated recombination and mosaicism.

Genes can circumvent knockout strategies and reinitiate transcription through various mechanisms such as exon skipping, nonsense-associated altered splicing, or the use of cryptic splice sites^45–48^. This highlights the importance of knocking out the entire gene rather than just the critical exons to prevent any unexpected mRNA transcripts^45,46^. Our investigation into the lengths of all protein-coding genes in mice (Supplemental Table 1; The data was sourced from Mouse Genome Informatics^49^) and the subsequent finding that the median length is 15.8 kb provided a valuable reference point (Fig. 3e). For a comprehensive examination of Cre-recombinase efficiency at inter-*loxP* distances of up to 15 kb, we designed three new strains at the *ROSA26* locus—*Rosa*-10 kb, *Rosa*-12 kb, and *Rosa*-15 kb. Because the recombination efficiency of the Sox2-cre strain was superior (Figs. 3b and 3d), we used this strain to investigate recombination efficiency at these three new inter-*loxP* distances. The data suggest that beyond an inter-*loxP* distance of 10 kb, recombination without mosaicism becomes less likely. This is evident from the Sox2-cre x *Rosa*-10 kb flox cross resulting in only 22% of offspring exhibiting recombination (Fig. 3f), and the Sox2-cre x *Rosa*-12 kb (Fig. 3g) and *Rosa*-15 kb (Fig. 3h) flox combinations showing no recombination, but 14% mosaicism at an inter-*loxP* distance of 12 kb.

Collectively, these findings provide valuable insights into the dynamics of Cre-mediated recombination and mosaicism relating to Cre driver strain. They underscore the superior recombination efficiency of the Sox2-cre strain at inter-*loxP* distances of up to 10 kb, while concurrently highlighting the influence of the genetic locus. We observed a notable decrease in recombination efficiency correlating with an increase in inter-*loxP* distance across all three Cre-driver strains. Considering these findings, we propose that researchers design floxed alleles with an inter-*loxP* distance of less than 4 kb to achieve complete recombination, and greater than 8 to 10 kb for mosaic recombination. This recommendation is based on the observation that over 73% of offspring from the Sox2-cre x *Rosa*-4 kb cross exhibited completely recombined alleles, and greater than 72% of offspring from the CMV-cre x *Rosa*-8 kb cross exhibited mosaicism. To further enhance recombination efficiency and mosaicism in F1 crosses, we advocate utilizing the Sox2-cre and either the Ella-cre or CMV-cre driver strains, respectively.

### Inter-*loxP* distance of 3 kb or less is optimal for efficient recombination when using mutant *loxP* sites

We next examined how changes in the distances between *loxP* sites in mutant *loxP* floxed alleles affected recombination efficiency and mosaicism. This was accomplished using five unique double-floxed inverted open reading frame (*DIO*) mouse strains, which contain wildtype *loxP* and mutant *loxP* (*lox2272*) each with different inter-*loxP* distances—1.1, 2.2, 5.3, 6.9, and 8.1 kb (Supplementary Fig. 3a). One of the significant challenges, especially when creating floxed *DIO* alleles with different inter-*loxP* distances at a fixed locus, has been the lack of a precise method for integrating DIO transgenes ranging from 1 kb to 8 kb. The currently available CRISPR/Cas9 technology is slow and presents difficulties due to the variability and inefficiency of CRISPR/Cas9-mediated homology-directed repair for inserting transgenes larger than 1-3 kb. Furthermore, the insertion of *loxP* sites using CRISPR/Cas9 introduces its own set of challenges^16^—the efficiency of generating conditional knockout alleles by simultaneously inserting right and left loxP sites via the delivery of 2 single guide RNAs (sgRNAs) and 2 single-stranded oligo DNA nucleotides (ssODNs) as donors is less than 1%. To address these issues, we utilized a high-efficiency Bxb1 recombinase system that we recently developed^40^ to individually integrate five constructs with different inter-*loxP* site lengths into the *ROSA26* locus (Fig. 4a). This approach allowed us to rapidly, efficiently, and uniformly generate five new *DIO* floxed strains.

**Figure 4.**
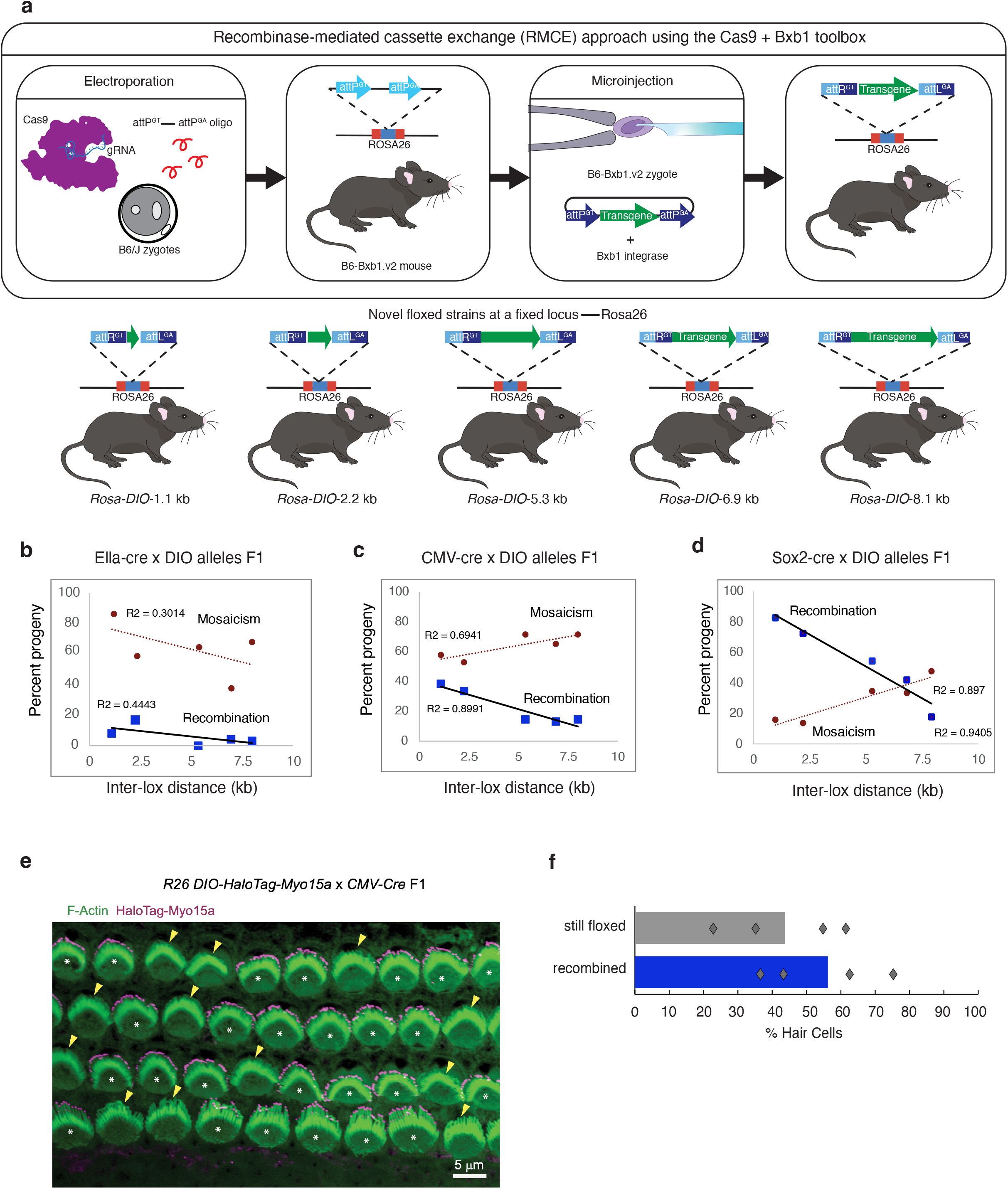
Impact of inter-*loxP* distances on Cre-mediated recombination and mosaicism in DIO mouse strains: from genotyping to single-cell analysis. (a) Study design for generating conditional alleles at a fixed locus (*ROSA26*) using the Cas9+Bxb1 toolbox. The Cas9+Bxb1 toolbox allows for accurate and efficient integration of large DNA constructs into a specific location, achieving higher efficiency compared to Cas9-mediated homology-directed repair (HDR). Bxb1 utilizes *attP* and *attB* attachment sites for DNA transgenesis. In our study, we strategically placed an attP attachment site within the *ROSA26* safe harbor locus of the B6/J mouse strain using CRISPR/Cas9-mediated HDR. This approach enabled us to seamlessly integrate DNA constructs with *DIO* alleles ranging from approximately 1 to 8 kb in size. Consequently, we successfully generated conditional knockout mice with single-copy transgenic modifications. (b, c, and d) Homozygous floxed male mice were mated with either female Ella-cre (b), CMV-cre (c), or Sox2-cre (d) mice, and the F1 offspring were genotyped using PCR from tail DNA to screen for floxed and recombined alleles. Offspring with complete recombination were categorized as recombined, those with no recombination as still floxed, and those with both recombined and floxed alleles as mosaic. The x-axis denotes the inter-*loxP* distance, and the y-axis denotes the percentage of recombination or mosaicism. The number of offspring genotyped to assess the percent of three possible outcomes: complete recombination, mosaicism, or no recombination are summarized in Supplementary Fig. 3b. (e) Mosaicism at the single-cell level: Confocal image of an 8-kb *Myo15a-Halo DIO* floxed allele x *CMV-*cre F1 cochlear sensory epithelium at postnatal day 2. Phalloidin stains filamentous actin (F-Actin), labeling the sensory hair cell cytoskeleton. TMR-conjugated *Halo Tag* ligand labels *Halo-Myo15a*, the product of recombination. *Halo-Myo15a* enriches at the tips of hair cell cytoskeletal projections. White asterisks indicate recombined cells expressing *Halo-Myo15a*. Yellow arrowheads indicate cells that escaped recombination, where the allele is still floxed. (f) Quantification of percentages from four different pups as points and the averages by bar plot.

Initially, we examined the impact of the Ella-cre driver strain on recombination and mosaicism across five mutant inter-*loxP* distances, ranging in length from 1.1 kb to 8.1 kb (Fig. 4b). For the offspring of the Ella-cre x *Rosa-DIO-*1.1 kb, an inter-*loxP* distance of 1.1 kb resulted in a mere 8% of the offspring showing recombination, while a noteworthy 88% displayed mosaic characteristics, and 4% remained floxed. Notably, the Ella-cre x *Rosa*-*-DIO-*2.2 kb cross led to a marginally better recombination rate of 16%, with 59% demonstrating mosaicism, and 23% remaining floxed. In the scenario of the Ella-cre x *Rosa*-*DIO*-5.3 kb cross, none of the offspring exhibited complete recombination, while 65% showed mosaicism, and 34% stayed floxed. For the Ella-cre x *Rosa*-*DIO*-6.9 kb pairing, 4% of the offspring showed full recombination, while 37% displayed mosaicism, and a 58% stayed floxed. Finally, in the Ella-cre x *Rosa*-*DIO*-8.1 kb *DIO* cross, 3% of the offspring showed full recombination, while a majority of 68% displayed mosaicism, and 28% stayed floxed (Fig. 4b).

Next, we delved into the impact of the CMV-cre-driver strain on recombination and mosaicism across five diverse mutant inter-*loxP* distances (Fig. 4c). With the *Rosa*-*DIO-*1.1 kb allele, the CMV-cre F1 progeny yielded a recombination rate of 38%, exhibited mosaicism in 57% of instances, and roughly 4% remained floxed. In the scenario of the *Rosa*-*DIO-*2.2 kb allele, 33% of the progeny demonstrated full recombination, 52% manifested mosaicism, and 13% remained floxed. In the case of the *Rosa-DIO-*5.3 kb allele, 14% of the progeny showed full recombination, a notable 71% displayed the mosaic state, and 14% remained floxed. For the *Rosa-DIO-*6.9 kb allele, 13% of the progeny exhibited full recombination, 65% displayed the mosaic state, and 21% remained floxed. Finally, for the *Rosa*-*DIO-*8 kb allele, 14% of the progeny demonstrated full recombination, a majority of 71% preserved the mosaic state, and 14% remained floxed (Fig. 4c).

Lastly, we investigated the impact of the Sox2-cre strain on recombination and mosaicism across five mutant inter-*loxP* distances (Fig. 4d). Upon analyzing the *Rosa*-*DIO-*1.1 kb allele, we observed that 81% of the Sox2-cre F1 progeny demonstrated recombination, 16% displayed mosaicism, and a small fraction of 2% remained without any recombination. When considering the *Rosa*-*DIO-*2.25 kb allele, we found that 71% of the progeny showed complete recombination, while 14% presented with the mosaic state, and an equivalent percentage remained floxed. In the case of the *Rosa*-*DIO-*5.3 kb allele, 53% of the progeny exhibited full recombination, 34% displayed the mosaic state, and 11% were floxed. Regarding the *Rosa*-*DIO-*6.9 kb allele, 41% of the progeny showed complete recombination, 33% showed a mosaic state, and 25% remained floxed. Finally, for the *Rosa-DIO-*8.1 kb allele, a mere 18% of the progeny demonstrated complete recombination, 47% were mosaic, and 34% remained floxed.

To investigate Cre-Lox mosaicism at the single-cell level, we used a cross with a high proportion of mosaic F1 progeny, CMV-cre x *Rosa*-*DIO-*8.1 kb. Upon recombination, the *Rosa-DIO-*8.1 kb allele induced the expression of the inner ear sensory cell-specific transcript Myo15a conjugated to the self-labeling HaloTag^50^. The neatly patterned single layer of cells in the cochlear sensory epithelium provides an ideal canvas for viewing cell-by-cell mosaicism. We isolated cochleae from F1 pups at postnatal day 2, labeled them with TMR-conjugated HaloTag ligand, and imaged the sensory epithelium. The mosaic pattern was striking, showing a clear variegation of sensory cells with and without HaloTag labeling at the distal cytoskeletal projections in a single field of view (Fig. 4e). Four different F1 mosaic offspring were analyzed, and the degree of mosaicism varied by individual, ranging from 37% to 76% of cells recombined. On average, a slight majority of cells were recombined, at 55% (Fig. 4f). Notably, tail-tip PCR-based genotyping of the CMV-cre x *Rosa-DIO*-8.1 kb F1 offspring (28 offspring from 4 litters) showed 71% mosaicism (Fig. 4c). This highlights the potential challenge of Cre-Lox configurations prone to mosaicism: experimental outcomes may be skewed due to the presence of unexpected mosaic individuals. Conversely, there may be cases in which having both recombined and still-floxed “escaper” cells within a single field of view can be advantageous, as cells that fail to recombine could act as internal controls for phenotypic comparisons of neighboring cells (*manuscript under review*).

Our experimental findings collectively indicate that an increase in the *DIO* inter-*loxP* distance from 1.1 to 8.1 kb is associated with a corresponding decrease in the percentage of recombined loci. It is noteworthy that the offspring of the Ella-cre x *DIO* cross exhibited the lowest recombination levels. Specifically, even with the 1.1 kb *DIO* allele, the percentage of recombined loci was less than 20%, and this percentage was even lower for all other tested Ella-cre x *DIO* progeny. Interestingly, no recombination was observed in the offspring of the Ella-cre x *DIO*-5.3 kb cross. However, among the three Cre-driver strains tested, Sox2-cre demonstrated superior recombination efficiency; the Sox2-cre x *DIO*-1.1 kb and Sox2-cre x *DIO-*2.2 kb crosses surpassed a recombination rate of 70%. The next tested distance (*DIO*-5.3 kb inter-*loxP* distance) resulted in a recombination rate of approximately 50%, and the percentage of mosaic loci steadily increased from 14% for *DIO*-2.2 kb to 34% for *DIO*-5.3 kb. This trend persisted when the mutant inter-*loxP* distance was extended from 5.3 kb to either 6.9 kb or 8.1 kb. These results underscore the complex relationship between the inter-*loxP* distance, mutant *loxP* sites, recombination efficiency, and the specific Cre-driver strain used in the study. Based on these observations, we suggest that researchers design *DIO* floxed alleles with an inter-*loxP* distance of less than 3 kb and ∼8 kb to maximize recombination efficiency versus mosaicism, respectively. This suggestion is based on the finding that over 71% of offspring from the Sox2-cre x *DIO*-2.2 kb cross exhibited completely recombined alleles and approximately half (47%) or more of offspring from all Cre x *DIO-8.1* kb cross exhibited mosaicism.

### Comparative analysis of wildtype and mutant *loxP* sites: impact on recombination efficiency and mosaicism

To more precisely evaluate the impact of using wildtype versus mutant loxP sites on recombination efficiency and mosaicism, we generated two distinct mouse strains, both with a fixed inter-*loxP* distance of 6.9 kb, but at different loci (*Rhbdf2* and *Rhbdf1*), with different *loxP* sites, either wildtype or mutant *lox71/66* sites (Fig. 5a). We selected these loci because of our prior experience with them as well as their relevance to human disease (see below). Our findings indicate that the use of wildtype *loxP* sites considerably improves recombination efficiency. This was evident when we compared the recombination efficiency of wildtype *loxP* (6.9 kb) and *lox71/66* (6.9 kb) using three different Cre-driver strains. The complete recombination efficiency improved from 4% with Ella-cre, to 13% with CMV-cre, and 42% with Sox2-cre strain when utilizing wildtype *loxP* sites. However, it is important to note that none of the three Cre-driver strains induced complete recombination in the mouse strain with mutant *loxP* sites (Fig. 5a). In a similar vein, when we compared wildtype *loxP* at the *Rhbdf2* locus (2.7 kb) and *lox71/66* at the *Rhbdf1* locus (2.9 kb) using the Sox2-cre strain, we observed a similar trend (Fig. 5a). There was an improvement in recombination efficiency from 42% to 72% when the inter-*loxP* distance was reduced from 6.9 to 2.7 kb while utilizing the wildtype *loxP* sites (Fig. 5a-b). Interestingly, we observed a lower recombination rate increase in Sox2-cre x 2.9 kb Mut *lox71/66* F1 progeny, reduced by 22 percentage points when compared to the Sox2-cre x 2.7 kb WT *loxP* F1 progeny. However, there was an increase from 0% to 50% in complete recombination rate in Sox2-cre x 2.9 kb *Mut lox71/66* F1 progeny compared with the Sox2-cre x 6.9 kb Mut *lox71/66* F1 progeny (Fig. 5c). These data suggest that when the inter-*loxP* distance was reduced from 6.9 kb to 2.9 kb, even when utilizing the mutant *loxP* sites, there was clear improvement in recombination.

**Figure 5.**
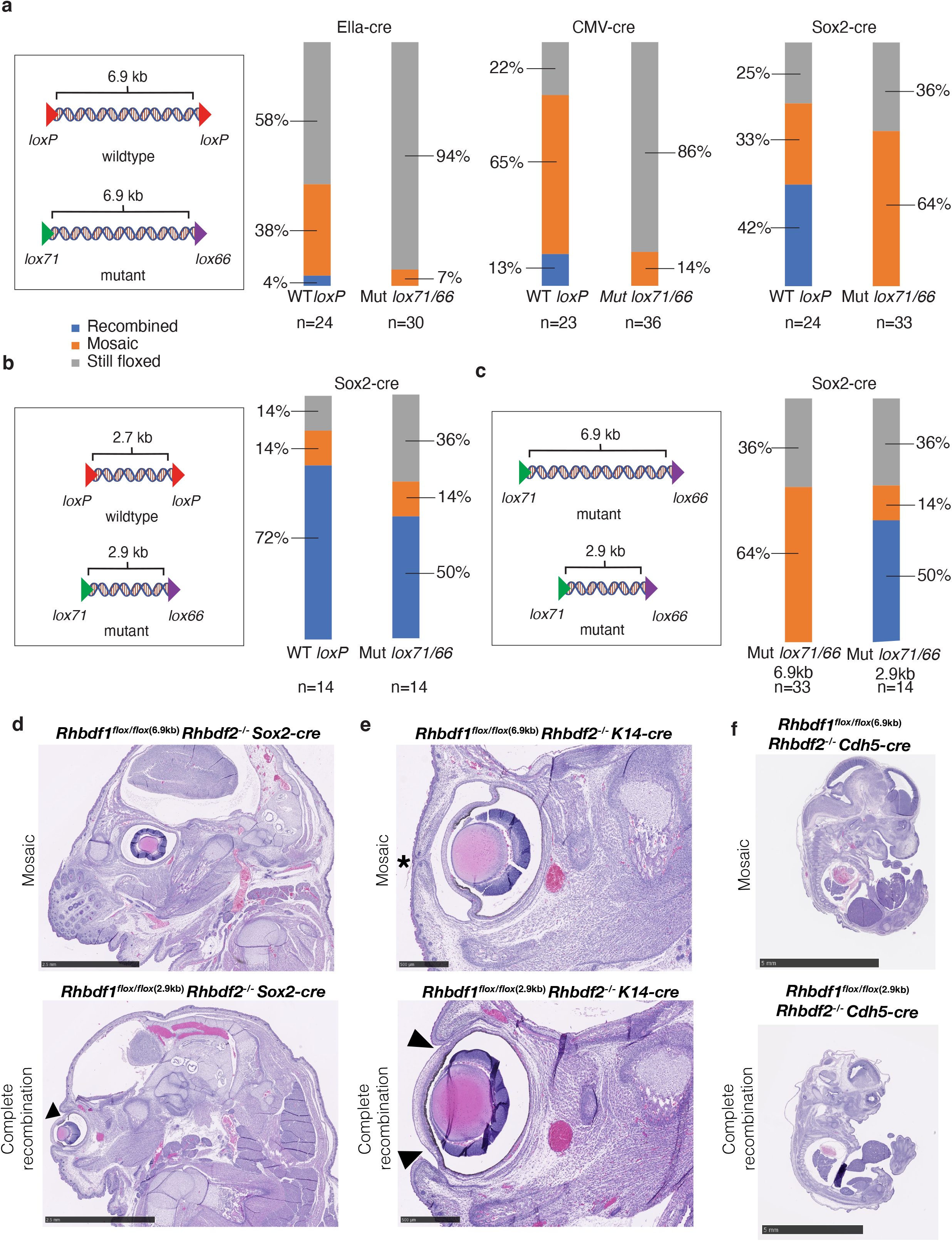
Inter-*loxP* distance and site type influence on Cre-mediated recombination efficiency and mosaicism. (**a**) Male homozygous floxed mouse strains, with a fixed inter-*loxP* distance of 6.9 kb at different loci and carrying either wildtype *loxP* sites or mutant *loxP* sites (*lox71* and *lox66*), were bred with female Ella-cre, CMV-cre, or Sox2-cre mice. (**b**) Male homozygous floxed mouse strains, with an inter-*loxP* distance of 2.7 kb and 2.9 kb at different loci, having wildtype *loxP* sites and mutant *lox71*/*66* sites, respectively, were bred with female Sox2-cre mice. (**c**) Male homozygous floxed mouse strains, with an inter-*loxP* distance of 6.9 kb or 2.9 kb at the same locus and with mutant *loxP* sites (*lox71* and *lox66*), were bred with female Sox2-cre mice. In Figures 5a, b, and c, F1 offspring were genotyped using PCR from tail DNA to detect floxed, mosaic, and recombined alleles. Offspring with both recombined and floxed alleles were considered mosaic. Note that percentages were rounded to the nearest whole number. (**d**) *Rhbdf1^-/-^ Rhbdf2^-/-^* double KO mice exhibit the ‘eyelids open at birth’ phenotype (indicated by arrows) and post-natal lethality^45^. However, when we generated *Rhbdf1* conditional KO mice with mutant *loxP* sites (*lox71* and *lox66*) and an inter-*loxP* distance of 6.9 kb (*Rhbdf1^flox/flox-^*^6.9k^*^b^*), and crossed with Sox2-cre mice, we were unable to generate *Rhbdf1^flox/flox-^*^6.9k^*^b^ Rhbdf2^-/-^* mice carrying the recombined *Rhbdf1* allele (top panel—mosaic). Conversely, reducing the inter-*loxP* distance of the *Rhbdf1^flox/flox-^*^6.9k^*^b^* conditional KO allele from 6.9 kb to 2.9 kb using CRISPR/Cas9-mediated deletion of up to 3 kb (*Rhbdf1^flox/flox-^*^2.9k^*^b^*), and crossing them with *Rhbdf2^-/-^* and Sox2-cre mice, resulted in the generation of the *Rhbdf1* recombined allele. This led to in *utero* lethality in *Rhbdf1^flox^*^2.9^*^/flox^*^2.9^ *Rhbdf2^-/-^* Sox2-cre mice (bottom panel—complete recombination). (**e**) The ‘eyelids open at birth’ phenotype (indicated by arrows) was observed in the *Rhbdf1^flox/flox^*^2.9k^*^b^ Rhbdf2^-/-^* K14-cre mice (bottom panel—complete recombination) but not in the keratinocyte-specific knock-out *Rhbdf1^flox/flox-^*^6.9k^*^b^ Rhbdf2^-/-^* K14-cre mice (top panel—mosaic). (**f**) *In utero* lethality was only observed with the *Rhbdf2^-/-^ Rhbdf1^flox/flox-^*^2.9k^*^b^* conditional KO allele (bottom panel—complete recombination), not with the *Rhbdf2^-/-^ Rhbdf1^flox/flox-^*^6.9k^*^b^* allele (top panel—Mosaic), when endothelial-specific Cdh5-cre was utilized.

To deepen our understanding of our research findings, we performed a thorough analysis of the distance between *loxP* sites using mutant *lox71/66* floxed alleles. The *Rhbdf1* gene, a member of the rhomboid family, has been associated with human cancer^51,52^. In previous studies, we engineered mice to be constitutively homozygous-null for *Rhbdf1* (*Rhbdf1^-/-^)*^45^. These mice displayed growth retardation, brain hemorrhage, cardiac hypertrophy, and unfortunately, did not survive beyond the 14th postnatal day^45^. To discern the specific role of *Rhbdf1* in various tissues, we engineered *Rhbdf1* conditional-ready mice (*Rhbdf1^flox/flox-^*^6.9k^*^b^*). This was accomplished by positioning the *lox71* site in intron 1-2 and the *lox66* site in exon 18, which are approximately 6.9 kb apart (Supplementary Fig. 4). Both mutant *loxP* sites were confirmed to be correctly targeted, with no unintended mutations, through PCR and Sanger sequencing. We first bred these *Rhbdf1^flox/flox-^*^6.9k^*^b^* mice with Ella-cre, CMV-cre, and Sox2-cre mice to produce tissue-wide homozygous-null mice. We anticipated that the deletion of *Rhbdf1* across all tissues would lead to multiple organ pathologies. However, the offspring from these crosses, where breeders were hemizygous for Sox2-cre (or homozygous for either Ella- or CMV-cre) and homozygous for the *Rhbdf1* floxed allele, were unexpectedly healthy, with no observable phenotypic or histological abnormalities (Supplementary Fig. 5). Further analysis utilizing PCR and probe-based assays revealed that despite the expression of the Cre transgene in all tested tissues, the recombinase failed to induce excision in most tissues. This resulted in mosaic recombination, leading to a viable phenotype. We hypothesized that the inter-*loxP* distance of 6.9 kb with the mutant *loxP* sites might be too large for efficient recombination at the *Rhbdf1* locus. To test this hypothesis, we used CRISPR/Cas9-mediated gene editing to reduce the inter-*loxP* distance from 6.9 kb to 2.9 kb. This was achieved by deleting non-critical exons 4 through 12, thereby creating *Rhbdf1^flox/flox-^*^2.9k^*^b^* mice (Supplementary Fig. 4). We have previously demonstrated that the deletion of these non-critical exons does not affect the viability or fertility of mice^45^. Finally, we crossed these *Rhbdf1^flox/flox-^*^2.9k^*^b^* mice with Sox2-cre driver strain mice. This resulted in the deletion of exons 2 through 18 via complete recombination and the expected phenotype—growth retardation, cardiac hypertrophy, and postnatal lethality (Supplementary Fig. 5b-d). Importantly, it appears that the *Rhbdf1* locus does not have stoichiometric limitations on Cre activity. This is based on our observation of inefficient Cre-recombination at other loci (*8.1-kb DIO floxed-Myo15a* allele with CMV-cre; Fig. 4e). Collectively, these findings suggest that 7 kb exceeds the allowable length for effective Cre-mediated recombination using these Cre driver strains at this locus.

The knockout of both *Rhbdf1* and *Rhbdf2* genes results in embryonic lethality^45^. Interestingly, when we attempted to replicate this phenotype using a combination of *Rhbdf1^flox/flox-^*^6.9k^*^b^* allele, Sox2-cre, and *Rhbdf2^-/-^* mice, we were unable to do so (Fig. 5d, upper panel). However, when we used the *Rhbdf1^flox/flox-^*^2.9k^*^b^* allele in combination with Sox2-cre and *Rhbdf2^-/-^* mice, the offspring exhibited embryonic lethality (Fig. 5d, lower panel). This finding supports our observation that for efficient Cre-recombination at the *Rhbdf1* locus, the optimal distance between mutant *loxP* sites is less than 3 kb. Furthermore, we discovered that the ‘open eyelid birth’ phenotype in *Rhbdf1* and *Rhbdf2* double knockout mice is regulated by the keratinocyte-specific expression of *Rhbdf1* and *Rhbdf2* (Fig. 5e, lower panel), as it is reproducible using a tissue-specific K14*-*cre driver. Again, lack of complete recombination in *Rhbdf1^flox/flox-^*^6.9k^*^b^*, K14-cre, *Rhbdf2^-/-^* mice, failed to show the ‘open eyelid birth’ phenotype with keratinocyte-specific knockout (Fig. 5e, upper panel). We also found that endothelial-specific expression is crucial for survival. This was evidenced by the observation that the endothelial-specific deletion of *Rhbdf1* in *Rhbdf2* knockout mice using the Cdh5-cre driver resulted in embryonic lethality (Fig. 5f). These findings provide valuable insights into the role of *Rhbdf1* and *Rhbdf2* in embryonic development and survival.

Overall, these findings shed light on the consequences of *Rhbdf1-KO* mosaicism, emphasizing the importance of considering inter-*loxP* distance in genetic studies, especially in the presence of mutant alleles.

### Impact of zygosity on recombination efficiency in Cre-Lox system

We next explored the impact of zygosity of the floxed alleles on recombination efficiency (Fig. 6a). We created a series of breeding schemes using Ella-cre (homozygous) or Sox2-cre (hemizygous) females crossed with either heterozygous or homozygous males from a variety of floxed strains. Strains tested included both wild-type and mutant *loxP* sites, both *ROSA26* and *Rhbdf1* genetic loci, and a range of inter-*loxP* distances from 0.8 kb to 8 kb. For heterozygous crosses, only offspring that inherited a parental floxed allele, regardless of recombination state, were considered. In the case of Ella-cre x *Rosa*-2.25 kb, we found that heterozygous offspring displayed 54% mosaicism and 38% recombination, while homozygous offspring exhibited 50% mosaicism and only 14% recombination (Fig. 6b). These results indicate that heterozygosity led to higher complete recombination efficiency in this breeding scheme. Our examination of other floxed alleles and the Sox2-cre driver strain yielded consistent results.

**Figure 6.**
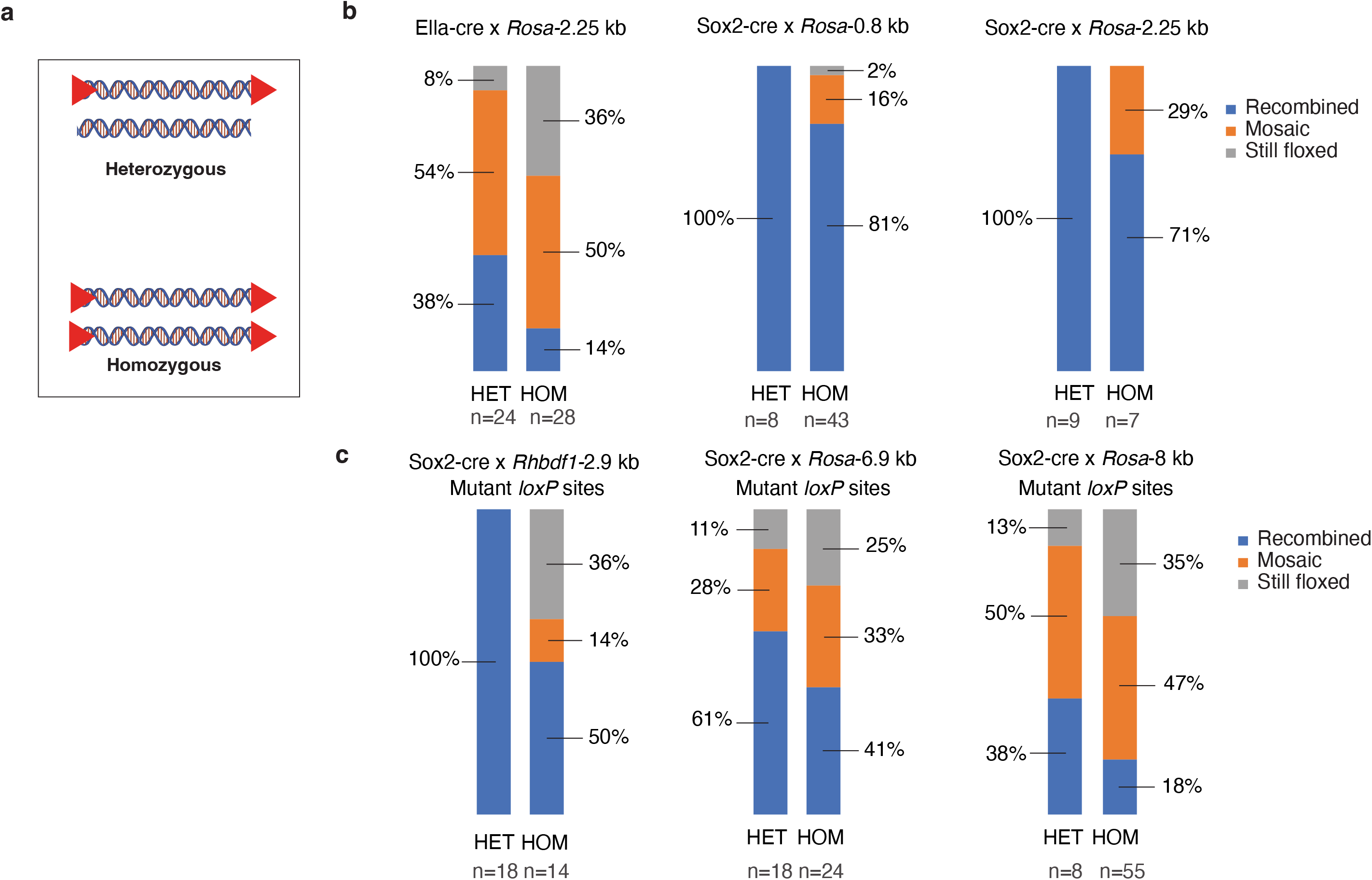
Impact of the zygosity of the floxed allele on recombination efficiency. (**a**) Schematic representation of the heterozygous (HET) versus homozygous (HOM) floxed alleles. (**b and c**) Male homozygous and heterozygous floxed mouse strains with either wildtype *loxP* (**b**) or mutant *loxP* (**c**)— *Rosa*-0.8 kb, *Rosa*-2.25 kb, *Rhbdf1*-2.9 kb, *Rosa*-6.9 kb, and *Rosa*-8 kb—were bred with female Ella-cre or Sox2-cre mice. The F1 offspring were then genotyped using PCR and categorized as recombined, still floxed, or mosaic. Note that the percentages were rounded to the nearest whole percent.

When examining Sox2-cre x *Rosa*-0.8 kb (wildtype *loxP* sites), Sox2-cre x *Rosa*-2.25 kb (wildtype *loxP* sites), and Sox2-cre x *Rhbdf1*-2.9 kb (mutant *loxP* sites), all instances of heterozygosity resulted in 100% recombination. However, in the case of homozygosity for Sox2-cre x *Rosa*-0.8 kb, only 81% recombination was achieved (with 16% mosaicism and 2% of the alleles remaining floxed). Similarly, for homozygous Sox2 -cre x *Rosa*-2.25 kb (wildtype *loxP* sites), only 71% underwent recombination (with 29% mosaicism). Lastly, in the scenario of homozygosity for Sox2-cre x *Rhbdf1*-2.9 kb (mutant *loxP* sites) only 50% experienced recombination (with 14% mosaicism and 36% still floxed) (Fig. 6c).

Heterozygosity was associated with higher recombination rates in crosses that were overall less efficient as well. The Sox2-cre x *Rosa-*6.9 kb and Sox2-cre x *Rosa*-8 kb crosses also favored heterozygosity. For *Rosa*-6.9 kb (mutant *loxP* sites), 61% of progeny were recombined (28% mosaic, 11% still floxed) when breeders were heterozygous versus only 41% recombined (33% mosaicism, 25% still floxed) when homozygous. Similarly, for *Rosa*-8 kb (mutant *loxP* sites), recombination efficiency was 38% (50% mosaic, 13% still floxed) for heterozygous breeders and only 18% (47% mosaic, 35% still floxed) for homozygotes (Fig. 6c).

In all breeding schemes, regardless of Cre driver strain, wildtype or mutant *loxP*, genomic locus or inter-*loxP* distance, heterozygous male breeders for the floxed allele yielded better recombination efficiency compared to homozygous male breeders. Collectively, these findings highlight the relationship between zygosity and recombination efficiency/mosaicism in our experimental setup.

### Impact of breeder age on recombination efficiency in Cre-Lox experiments

The age of Cre-driver strain breeders is another variable with potential impacts on recombination efficiency. To investigate this, we used female CMV-cre and Sox2-cre mice and collected litters from breeders ranging in age from 5 to 38 weeks old at the time of mating (Fig. 7a). Furthermore, we tested four different floxed alleles with inter-*loxP* distances ranging from 0.8 kb to 8 kb. We determined the percentage of recombined pups from a total of 4-8 litters for each floxed strain.

**Figure 7.**
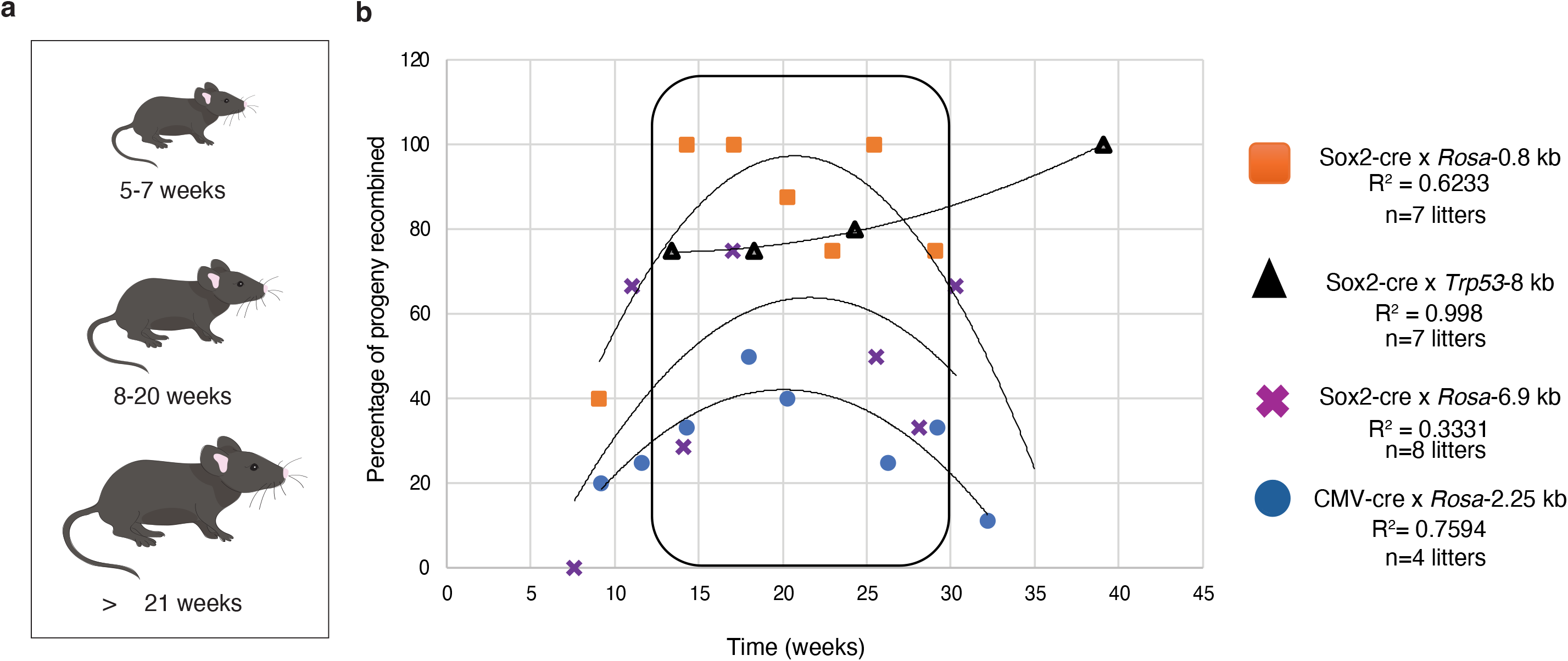
Age-dependent recombination efficiency in mouse breeders and its impact on Cre-mediated recombination of floxed alleles. (**a**) Representation of mouse breeders ranging in age from 5 to 38 weeks was used to determine recombination efficiency. (**b**) Effects of age on Cre-mediated recombination of four different floxed alleles. Each data point represents the percentage of recombination in a litter of pups, coded by cross of origin, as labelled on plot. X-axis is the age (in weeks) of the Cre-driver breeder at the time of progeny birth.

Recombination efficiency varied between litters for all four floxed alleles. In general, it was lower in litters derived from very young Cre breeders (less than 10 weeks of age at the time of birth) and older breeders (more than 25 weeks). To determine whether breeder age was truly correlated with recombination rate, we fitted a non-linear curve to each floxed allele data set. The correlation was strongest (R² = 0.998) for the Sox2-cre x *Trp53*-8 kb allele, with a slightly positive, nearly linear relationship between Cre breeder age and the percentage of recombined progeny. However, this trend was an exception, as the other three alleles each fit best to a downward-opening parabolic curve (R² = 0.759, 0.333, and 0.623, respectively for CMV-cre x *Rosa*-2.25 kb, Sox2-cre x *Rosa*-6.9 kb, and Sox2-cre x *Rosa*-0.8 kb), indicating an ideal breeder age in the median of the range, with a consistent consensus of approximately 20 weeks yielding peak efficiency. For three of the four floxed strains tested, litters collected from Cre breeders older than 20 weeks faced a decline in recombination efficiency. This decline was not observed in the Sox2-cre x *Trp53*-8 kb litters, which may be attributed to a lack of data points for advanced age (more than 20 weeks) Cre breeders compared to the other floxed strains measured (Fig. 7b).

We therefore conclude that the age of the Cre-driver strain at the time of breeding should be an additional consideration in Cre-Lox experimental design. This data corroborates previous, limited evidence supporting age-based effects in Cre-driver mouse strains^53^. For optimal recombination rate, we recommend using Cre breeders that are at least 8 weeks old, but no older than 20 weeks.

### Efficiency of Cre mediated recombination across different genomic loci in conditional knockout mouse strains

A majority of our studies herein focused on *Rosa* floxed alleles, which have the benefit of consistency with respect to genomic loci. However, studies involving Cre-Lox for the purpose of conditional knockout, for example, will require Cre recombination targeted to other regions of the genome. Genomic locus should therefore be considered as an additional variable affecting recombination rate. To address this question, we used the Sox2 - cre strain crossed with twelve different conditional knockout mouse strains with floxed alleles across different regions of the mouse genome (Fig. 8 and Supplementary Figs. 6-13). This panel spanned a total of nine autosomal chromosomes. All conditional knockout alleles had inter-*loxP* distances <4 kb to support optimal recombination, with a range of 0.8 kb to 3 kb.

**Figure 8.**
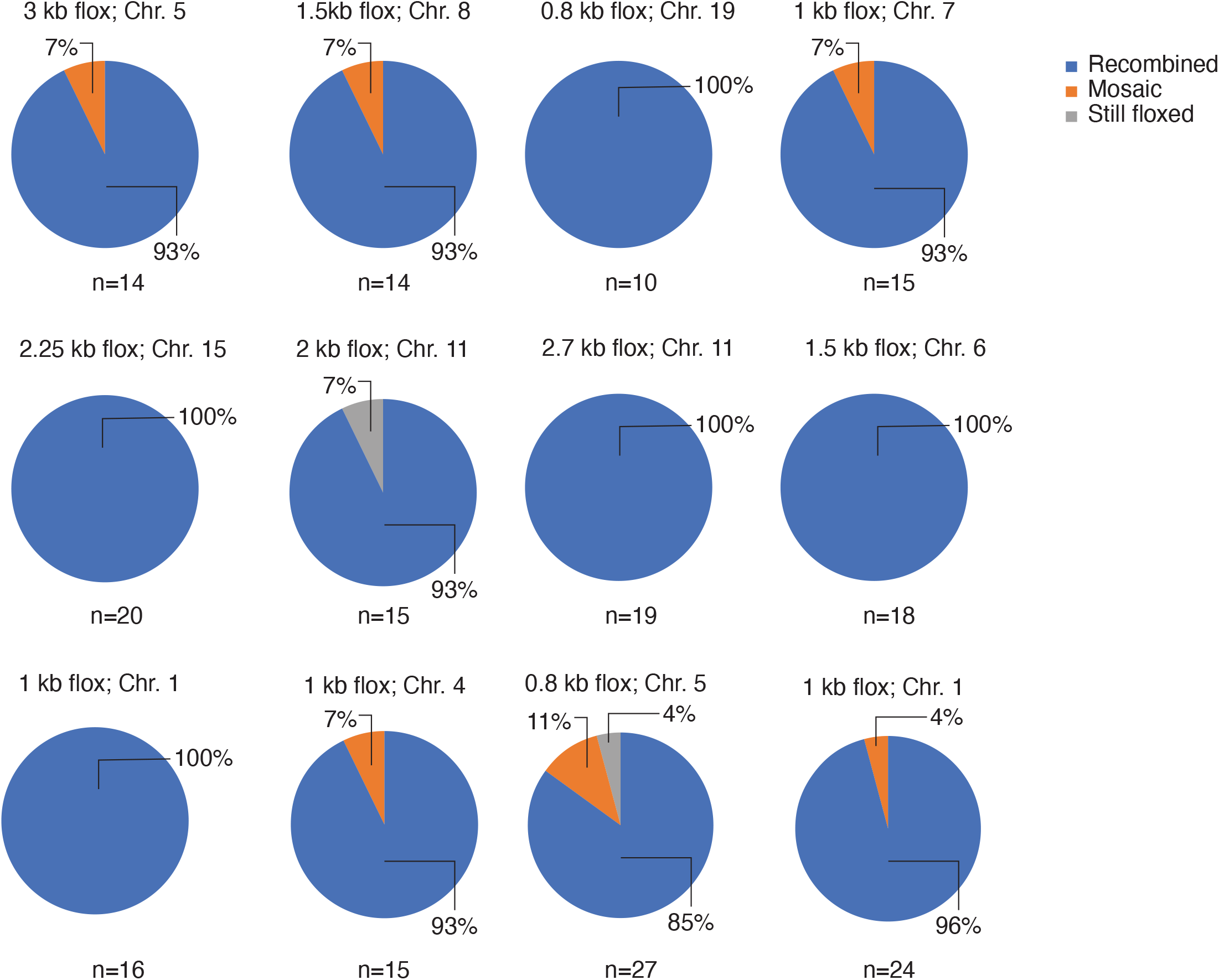
Genome-wide characterization of homozygous conditional knockout mouse strains using Sox2-cre recombination. Twelve different male homozygous conditional KO strains, with inter-*loxP* distances ranging from 0.8 to 3 kb at various loci across the mouse genome, were crossed with the Sox2-cre strain. The F1 offspring were then genotyped to identify floxed, mosaic, and recombined alleles. Note that the percentages were rounded to the nearest whole percent.

Irrespective of chromosomal location, recombination efficiency was high for each cross, ranging from 85% to 100% recombined F1 offspring. Rates of mosaicism in offspring were low, ranging from 0% to 11% of offspring, and failure to recombine was rare, ranging from 0% to 7% still floxed. Five of the twelve crosses showed 100% recombination, with floxed alleles on chromosomes 1, 6, 11, 15, and 19. Mosaicism was observed in offspring from crosses with floxed alleles on chromosomes 1, 4, 5, 7, and 8. Only two crosses yielded litters with any non-recombined progeny: Sox2-cre x Chr.11 2 kb flox and Sox2-cre x Chr.5 0.8 kb flox. Thus, chromosome location does not appear to be a strong predictor of recombination efficiency in this study. It is informative to compare the crosses with floxed alleles on the same chromosome, for example Sox2-cre x Chr.11 2 kb flox (93% recombined) vs. Chr.11 2.7 kb flox (100% recombined), or Chr.5 0.8 kb flox (85% recombined) vs. Chr.5 3 kb flox (93% recombined). These cases demonstrate that the efficiency of recombination can indeed vary. This variation is minor, however, and is based on the genomic locus, even when considering loci on the same chromosome (Fig. 8).

Therefore, we conclude from this panel that conditional knockout alleles with an inter-*loxP* distance <4 kb should have high recombination efficiency regardless of the floxed allele’s location in the mouse genome.

## Discussion

The flexibility of the Cre-Lox system extends beyond its standalone capabilities. It can be seamlessly integrated with other gene-editing systems, such as CRISPR-Cas9 and Cas12a, thereby expanding its potential applications and empowering researchers with a toolkit for genetic manipulation. Over the years, the Cre-Lox system has been employed in various groundbreaking studies. For instance, it played a crucial role in the development of techniques for identifying specific cell types, such as Brainbow, which enables the labeling and distinction of individual neurons, advancing our understanding of complex neural networks^12,29^. Notably, it was instrumental in generating a mouse strain for studying COVID-19^54^ and played a role in the development of vaccines for avian influenza^55^.

Despite the utility of the Cre-Lox system for genetic engineering, the use of Cre is associated with several limitations. First, Cre expression can be unpredictable and “leaky,” potentially interfering with the intended results. For example, the unexpected ectopic expression of Cre can lead to unanticipated recombination events during development. Second, inducible Cre expression may vary across different tissue types, with common inducers, like tamoxifen, exhibiting tissue-specific variations. Third, high levels of Cre expression have been associated with toxicity, although tamoxifen has also been implicated as a potential factor in the case of conditional *CreER^T^* drivers. In addition, efficient use of Cre-Lox is hindered by factors not specific to Cre expression. For example, not all tissue-specific promoters are truly specific, which can result in gene expression in unintended cell populations. In addition, differences among genes in their propensity for recombination may be influenced by the physical structure of chromatin. These previously documented issues collectively contribute to unexpected phenotypic outcomes, posing a significant obstacle to the generation of faithful mouse models of human diseases.

The findings we present here provide guidance to help researchers use Cre-Lox technology as effectively as possible in the face of the above limitations. With respect to the choice of a cre-driver strain, we show that Sox2-cre is more effective than Ella-cre or CMV-cre in achieving recombination. Furthermore, by examining varying inter-*loxP* distances (0.8 kb, 4 kb, 8 kb, 10 kb, 12 kb, and 15 kb), our work sheds light on how these distances impact recombination rates and mosaicism, aiding in the design of future experiments involving Cre-mediated recombination. Moreover, our findings suggest that tail DNA lysate could serve as a reliable predictor of recombination/mosaic patterns in the entire organism, streamlining the process of analyzing recombination and mosaicism in genetic studies.

Our results show that an inter-*loxP* distance of less than 3 kb is optimal for recombination when utilizing mutant *loxP* sites. To investigate the effects of mutant *loxP* sites on recombination efficiency, with a special emphasis on the role of varying inter-*loxP* distances in mutant *loxP* floxed alleles, we employed a high-efficiency Bxb1 recombinase system to incorporate five constructs, each with different inter-*loxP* site lengths, into the *Rosa26* locus. This strategy expedited the creation of five novel double-floxed inverted open reading frame (*DIO*) mouse strains. Our observations indicate that recombination efficiency diminishes as the inter-*loxP* distance extends from 1 kb to 8 kb. This insight is pivotal for the design of genetic experiments that involve Cre-mediated recombination. Our research also unveils that different Cre-driver strains, namely Ella-cre, CMV-cre, and Sox2-cre, have diverse impacts on recombination and mosaicism. For example, the progeny from the Ella-cre x *DIO* crosses demonstrated the lowest levels of recombination, with less than 20% recombination even with the 1 .1 kb *DIO* allele. In contrast, crosses involving the Sox2-cre driver strain exceeded a recombination rate of 70% with inter-*loxP* distances of 1 kb and 2.25 kb. This suggests that *DIO* floxed alleles with inter-*loxP* distances of less than 3 kb enable efficient recombination. Moreover, the research uncovers intriguing patterns in mosaicism. As the inter-*loxP* distance expands, the percentage of mosaic individuals also escalates, implying that larger inter-*loxP* distances may result in higher incidences of mosaicism. This could be a significant factor to consider when designing floxed alleles with mutant *loxP* sites.

Our research also explored the relationship between the age of Cre-driver strain breeders and the degree of mosaicism in their offspring, revealing that breeders aged between 10 and 20 weeks were most likely to achieve successful recombination. We embarked on this investigation by employing CMV-cre and Sox2-cre strains, collecting litters from breeders whose ages ranged from 5 to 38 weeks at the time of mating. We tested four distinct floxed alleles, with inter-lox distances spanning from 0.8 kb to 8 kb. We observed that recombination efficiency varied across litters for all four floxed alleles, with a general trend of lower efficiency in litters from extremely young breeders (under 10 weeks) and older breeders (over 25 weeks). Notably, for three out of the four floxed strains tested, litters from Cre breeders older than 20 weeks exhibited a decline in recombination efficiency. These findings underscore the importance of considering the age of the Cre-driver strain at the time of breeding when designing Cre-Lox experiments. This may be a concern in experimental designs that use multiple litters from the same breeding pair over time as replicates, where temporal changes in recombination efficiency may produce variable results.

We also investigated the influence of zygosity on the efficiency of recombination in floxed alleles. A series of breeding schemes was established, involving both homozygous and heterozygous crosses with a variety of floxed strains. The strains tested encompassed both wildtype and mutant *loxP* sites, different genetic loci (*ROSA26* and *Rhbdf1*), and a range of inter-*loxP* distances. Our findings indicate that heterozygosity consistently led to higher recombination efficiency across all tested parameters. This was observed even in less efficient crosses, suggesting a robust relationship between zygosity and recombination efficiency. For best reproducibility, parental zygosity should be consistent in experiments using multiple litters.

We examined the impact of using wildtype versus mutant *loxP* sites on recombination efficiency and mosaicism in mouse models. In particular, our findings from two distinct mouse strains, one with wildtype *loxP* sites and the other with mutant *lox71/lox66* sites, suggest that the use of wildtype *loxP* sites improves recombination efficiency. We found that when comparing the efficiency of wildtype *loxP* and *lox71/lox66,* the recombination efficiency for a 6.9 kb inter-*loxP* allele improved from 0% to 4%, 13%, or 42% depending on the Cre-driver strain. Further analysis was performed to understand the impact of inter-*loxP* distance on recombination efficiency. By reducing the inter-*loxP* distance from 6.9 kb to 2.9 kb, even when utilizing mutant *loxP* sites, there was a dramatic increase in recombination, rising from no recombination to 50%. We also explored the specific role of the *Rhbdf1* gene in various tissues. *Rhbdf1* conditional-ready mice were generated by positioning *lox71* and *lox66* sites approximately 6.9 kb apart. Surprisingly, homozygous-null mice resulting from crosses with Cre-driver strains did not display expected phenotypic abnormalities, indicating inefficient recombination. To address this, the inter-*loxP* distance was reduced to 2.9 kb using CRISPR/Cas9-mediated gene editing. This modification resulted in complete recombination and the expected phenotype in offspring. Crucially, our findings suggest that for efficient Cre-recombination, the optimal distance between mutant *loxP* sites is less than 3 kb. This conclusion was supported by experiments attempting to replicate the embryonic lethality phenotype seen with the knockout of both *Rhbdf1* and *Rhbdf2* genes. Overall, our findings underscore the importance of considering inter-*loxP* distance in genetic studies, especially when dealing with mutant floxed alleles.

Our research, while primarily centered on an array of *ROSA26* floxed alleles, also encompasses a broader genomic scope. In this context, we employed the Sox2-cre strain in conjunction with twelve distinct conditional knockout mouse strains. These strains possessed floxed alleles with inter-*loxP* distances of <4 kb, scattered across various regions of the mouse genome. The outcomes were promising, with all crosses demonstrating a high recombination efficiency, ranging from 85% to 100%, regardless of their chromosomal positions. Furthermore, the instances of mosaicism in offspring were minimal, and instances where recombination failed to occur were exceedingly rare. These findings lead us to propose that, irrespective of their genomic location, conditional knockout alleles with an inter-*loxP* distance of <4 kb are likely to exhibit high recombination efficiency. This insight holds significant implications for genetic manipulation in mouse models, and is encouraging for studies involving conditional knockout, for example, where the locus of the floxed allele cannot be controlled.

In conclusion, our study systematically optimized Cre-mediated recombination and efficiency in mice and provides guidelines for efficient design of floxed alleles with wildtype and mutant *loxP* sites. Additionally, we provide guidelines for optimal inter-*loxP* distance for efficient recombination. Our findings underscore the complex interplay of factors such as inter-*loxP* distance, Cre driver strain, breeder age, and zygosity on recombination efficiency. This highlights the necessity of fine-tuning experimental conditions to yield robust and reproducible results. Key findings of our research include the exceptional performance of the Sox2-cre strain and an optimal inter-*loxP* distance of less than 4 kb when using wildtype *loxP* sites for recombination. We discovered that a reduction in the inter-*loxP* distance from 6.9 kb to 2.9 kb enhanced recombination, even when using mutant *loxP* sites. This suggests that for efficient Cre-recombination, the optimal distance between mutant *loxP* sites is less than 3 kb. Expanding beyond *ROSA26* floxed alleles, our study explores a wider genomic landscape, demonstrating that high recombination efficiency can be achieved across diverse genomic locations, provided the inter-*loxP* distance is <4 kb. These collective findings deepen our understanding of Cre-mediated recombination, providing a solid foundation for designing future genetic experiments.

Our research opens up new avenues for generating mosaic animals characterized by sporadic gene activation. Producing a mosaic of recombined and still-floxed “escaper” cells can be very convenient for analysis of cellular features at the single-cell or tissue level. For example, we recently used non-recombined sensory cells in the cochlea as internal controls for neighboring conditional knockout cells while measuring the length of cytoskeletal projections (*manuscript under review*). This ensured that our comparisons were as accurate as possible, given that cellular features change depending on cochlear position. Measurements comparing directly adjacent cells are only possible with mosaic recombination.

Mosaic recombination also has a unique advantage for cancer research. It is worth noting that there is a fundamental difference between a transgenic model, where an oncoprotein is expressed in every cell of a tissue or cell lineage, and human cancers, where this is not the case. These mosaic animals could prove to be invaluable tools for simulating human cancer, particularly when germline embryonic transgene expression results in lethality^56^. For example, the progeny of K-*ras* floxed mice and Protamine-cre mice (which is active in haploid sperm) exhibit embryonic lethality due to the expression of the K-Ras oncoprotein during early development^57^. To overcome this issue and establish a tumorigenesis model using sporadic K-*ras* activation (instead of widespread activation), Guerra et al. produced offspring by crossing K-*ras* floxed mice with CMV-cre mice^58^. This resulted in a significant number of offspring reaching adulthood, presumably due to incomplete CMV-cre mediated recombination of the K-*ras* floxed allele. Furthermore, signs of pathology, such as respiratory problems, were observed when the progeny reached seven to eight months old^58^, suggesting that mosaicism can reduce the tumor burden. Our systematic analysis of different inter-*loxP* distances, different types of *loxP* sites, the age of the breeder strains, and the zygosity of the floxed alleles, should facilitate the creation of better mouse models of cancer. Notably, as reported by Guerra et al., reduction in tumor burden due to mosaicism could potentially prolong the lifespan of the mice, enabling longer-term studies. The knowledge gained here, along with alterations in the number of adeno-associated virus particles delivered, an increase in the wide availability of AAV-cre, and the combination of AAV serotype and promoter specificity, can help to selectively activate or silence conditional alleles, thereby generating non-germline mouse models to enable the study of cancer at single-cell resolution.

## Methods

### Mice

All the mouse strains listed in Table 1, as well as other strains such as *C57BL/6J-Gt(ROSA)26Sor^em7Mvw^/MvwJ (Bxb1.v2*)*, B6N(Cg)-Rhbdf1 tm1.1(KOMP)Vlcg/J, B6N:B6J-Rhbdf2<tm1a(KOMP)Wtsi>/Vhr,* B6.Cg-*Edil3^Tg^*(Sox2– cre)*^1Amc^*/J, B6.C-Tg(CMV-cre)1Cgn/J, and B6.FVB-Tg(EIIa-cre)C5379Lmgd/J were bred and maintained in a barrier facility at JAX with a 12-h light to 12-h dark cycle. All animals were maintained in accordance with institutional policies governing the ethical care and use of animals in research, under approved protocols. The strains listed in Table 1, denoted by an asterisk, were generated using Bxb1-mediated recombination-mediated cassette exchange (RMCE), as described previously^40^. Briefly, using a single 18.5 kb prototype construct, we generated five constructs that share sequence homology. Each construct includes *attB-GT* and *attB-GA* sites that have a matching cognate *attP-GT* and *attP-GA* sites in the *ROSA26* locus in the *Bxb1.v2* mice for Bxb1-mediated RMCE (Fig. 4a). Real-time PCR probes for tdTomato, YFP, and EYFP were designed to genotype mice using Transgenic Genotyping Services at JAX and Transnetyx. Additionally, they all have a fixed *loxP* site at the 5’ end and a flanking second *loxP* site. The five constructs differ in the length of the internal sequence, having either 4, 8, 10, 12, or 15 kb between *loxP* sites (Supplemental Fig. 14). To generate the *Rhbdf1* conditional mice (*Rhbdf1^flox/flox^* 6.9 kb), fertilized oocytes from C57BL/6J mice (stock #664) were electroporated using three sgRNAs (sg2472, sg2540, and sg2541) targeting intron 1–2 (*lox71)* and a site after exon 18 (*lox66*), respectively. For the left *lox71* insertion, sgRNA 2472: GTGCTCACAGCACCCGATAG and the ssODN:GAGTAGGTTTATACAGTGTTTTACCCTAGATTCCTATTGTTGGAGGATCTGTGGAGTTCAGGCCCAGAGCATGCCAGCCTCTATACCGTTCGTATAGCATACATTATACGAAGTTATTCGGGTGCTGTGAGCACCTCTGTAGGGCTTGCAATGCTTCTTCTAGGTTTAGGCCCATGGTGTCCCTGGAGACCTGTCAGGGT were used. For the right *lox66* insertion, the following sgRNAs (sg2540 and sg2541): GTTCTCAACCTGCCTGTG and GGGTCACGCCGCACAGGC, and the ssODN: TCACTGTTTCCTGATTTTTGGCTGGGGTCACGGGGGTA GGCCTTGGACATACAGTTGGACAAGAGAGTGGTTCTCAACCTGCCATAACTTCGTATAGCATACATTATACG AACGGTAGTGCGGCGTGACCCCTCTGGCAAACCTCTAGCTCCAGGATGGCTCAAAACAGTAGCAAAAG TCGCAACAAACTGTGTTAAAGG were used. Each sgRNA was delivered at a concentration of 150 ng/μl, along with Cas9 protein (PNABio) at 250 ng/μl. To generate the *Rhbdf1* conditional mice (*Rhbdf1^flox/flox-^*^2.9k^*^b^*) mice, fertilized oocytes from the homozygous *Rhbdf1* conditional mice (*Rhbdf1^flox/flox-^*^6.9k^*^b^*) were electroporated using two sgRNAs targeting intron 3-4 (sgRNA1020) and intron 12-13 (sgRNA 1021), respectively, to delete a 3 kb region between the two sgRNA targeting sites.

### Zygote electroporation or microinjection

The process of gene editing was performed using two distinct methods: microinjection (large DNA transgenesis using Bxb1 mRNA) or electroporation (CRISPR/Cas9-mediated gene deletion). The procedures for both electroporation and microinjection were carried out according to previously published protocols^40,42,59^. Briefly, zygotes were harvested from C57BL/6J females that had undergone superovulation and mating. These zygotes were then placed in a specially prepared embryo collection medium. Following this, the zygotes were either immediately subjected to manipulation or kept in an incubator at a temperature of 37 °C with 5% CO2 until the time of manipulation and transfer. After the manipulation process, the embryos were transferred into pseudopregnant recipients.

### Genotyping

Pregnant dams were euthanized by CO_2_ asphyxiation, performed in a manner consistent with the 2013 recommendations of the American Veterinary Medical Association Guidelines on Euthanasia. Offspring from Cre-driver x floxed strains were collected from pregnant mice at E17.5. Pups P1 to P3 were decapitated. Tail tips, no longer than 2 mm, were excised for DNA preparation. Tail tips were either stored overnight at 4°C or directly placed in 100 μl of lysis buffer containing 1.5 μl of Proteinase K, then incubated overnight at 55°C. Following this, they were denatured at 95°C for 5 minutes. Tail lysate was diluted either 1:4 (for adult mice) or 1:10 (for embryos or pups) in nuclease-free water, and 1 μl of the diluted lysate was used in a 20 μl PCR reaction. Where indicated, real-time PCR probes were designed to genotype mice or tissues using Transgenic Genotyping Services at JAX and Transnetyx. The list of primers used in this study is available in Supplemental Table 2.

### Cochlear Tissue Preparation and Fluorescence Microscopy

The inner ears were extracted from neonatal mice at postnatal day 2, and the cochlear sensory epithelium was isolated and exposed via dissection in PBS. Cochlea were fixed for 15 minutes in 4% paraformaldehyde (PFA, 15710, Electron Microscopy Sciences) at room temperature, then permeabilized in PBS with 0.5% Triton X-100. Samples were incubated in 0.5 µM TMR-conjugated HaloTag ligand (G8252, Promega) in PBS for 1 hour at room temperature, then incubated in FITC-conjugated phalloidin (P5282, Sigma) in PBS overnight at 4°C. Samples were washed 4 times in PBS with 0.05% Triton X-100, then post-fixed for 1 hour in 4% PFA at room temperature. Whole-mounts were prepared in 10% Mowiol (475904, EMD/Millipore; in 0.1 M tris-Cl, pH 8.5) on positively charged slides (M1021, Denville) with 18 x 18 mm No 1.5 coverslips (48366-045, VWR). Images were collected with Zeiss LSM 800 confocal laser-scanning microscope with a 63x NA1.4 oil immersion objective using the ZEN 2.6 software (Carl Zeiss AG) and enhanced in Adobe Photoshop CC 2023. For measurements in Figure 3F, recombined cells were identified by the enrichment of TMR HaloTag ligand signal at the tips of hair cell cytoskeletal projections. Quantities of recombined and still floxed cells were counted in ImageJ using the Cell Counter plugin.

### Bxb1-mediated insertion of *DIO* alleles into the *ROSA26* of the Bxb1.v2 mice

For a step-by-step protocol on plasmid donor DNA construction with attachment sites, zygote microinjection, embryo transfer, and RMCE, please see Erhardt et al. 2024. The B6. Bxb1.v2 mouse strain has pre-placed dual heterologous attachment sites attP-GT and attP-GA; The attP-GT sequence: 5’ GGTTTGTCTGGTCAAC CACCGCG<u>GT</u>CTCAGTGGTGTACGGTACAAACC 3’ and the attP-GA sequence: 5’ GGTTTGTCTGGTCAAC CACCGCG<u>GA</u>CTCAGTGGTGTACGGTACAAACC 3’. The donor plasmids have the complementary attachment sites attB-GA and attB-GT; The attB-GT sequence: 5’ ggcttgtcgacgacggcg<u>gt</u>ctccgtcgtcaggatcat 3’ and the attB-GA sequence: 5’ ggcttgtcgacgacggcg<u>ga</u>ctccgtcgtcaggatcat 3’. Successful RMCE resulted in the attR-GT and attL-GA sites: The attR-GT sequence: 5’ GGTTTGTCTGGTCAACCACCGCGGTctccgtcgtcaggatcat 3’ and the attL-GA sequence: 5’ ggcttgtcgacgacggcg<u>GA</u>CTCAGTGGTGTACGGTACAAACC 3’. Briefly, homozygous Bxb1.v2 mouse zygotes were microinjected with 30 ng/μl of donor DNA and 120 ng/μl of Bxb1 mRNA. The microinjection reagents were combined in nuclease-free IDT TE pH 7.5 (Integrated DNA Technologies; Tris 10 mM, EDTA 0.1 mM) supplemented with Rnasin (Promega) at 0.2 U/μl. A comprehensive, step-by-step screening method was used to screen founder animals using established protocols^40,59^ to identify transgene integration. This involved at least three PCRs: 1) transgene-specific PCR, 2) left RMCE PCR, 3) right RMCE PCR. PCR products were sequence-verified by Sanger sequencing.

### Histology

Performed as described previously^60^. Briefly, pregnant dams were euthanized by CO_2_ asphyxiation. Following CO_2_ asphyxiation, embryos were washed in ice-cold phosphate-buffered saline. The embryos (E15.5 to E17.5) were then preserved overnight in 10% neutral buffered formalin before being moved to a 70% ethanol solution. Subsequently, the embryos were paraffin embedded, sectioned in the mid sagittal plane, and stained with hematoxylin and eosin.

### Ethics statement

All the methods used in this study were approved by The Jackson Laboratory’s Institutional Animal Care and Use Committee (AUS # 22022 and # 11006). The mice involved in the study were kept at The Jackson Laboratory, located in Bar Harbor, ME, USA, and all institutional protocols and the Guide for the Care and Use of Laboratory Animals were strictly followed. The data and experiments presented comply with the ARRIVE guidelines, where relevant.

## Acknowledgements

We gratefully acknowledge the contribution of Scientific Services at JAX for their assistance with microinjections (Peter M. Kutny, Genetic Engineering Technologies) and histopathology, Stephen B. Sampson for critically reading the manuscript, and Gloria Fuentes for helping with the figures.

## Author contributions

Conceptualization and Supervision: B.T and V.H.; Performed the experiments: V.E., E.H., H.M., K.L., and B.T, Writing - original draft: E.H. and V.H.; Writing - review & editing: E.H., K.L., B.T., and V.H.; Funding acquisition: B.T. and V.H.

