## Supplemental Figure for "Large-Scale Genome-Wide Optimization and Prediction of the Cre Recombinase System for Precise Genome Manipulation in Mice"

### Supplemental Figure 1

**a**

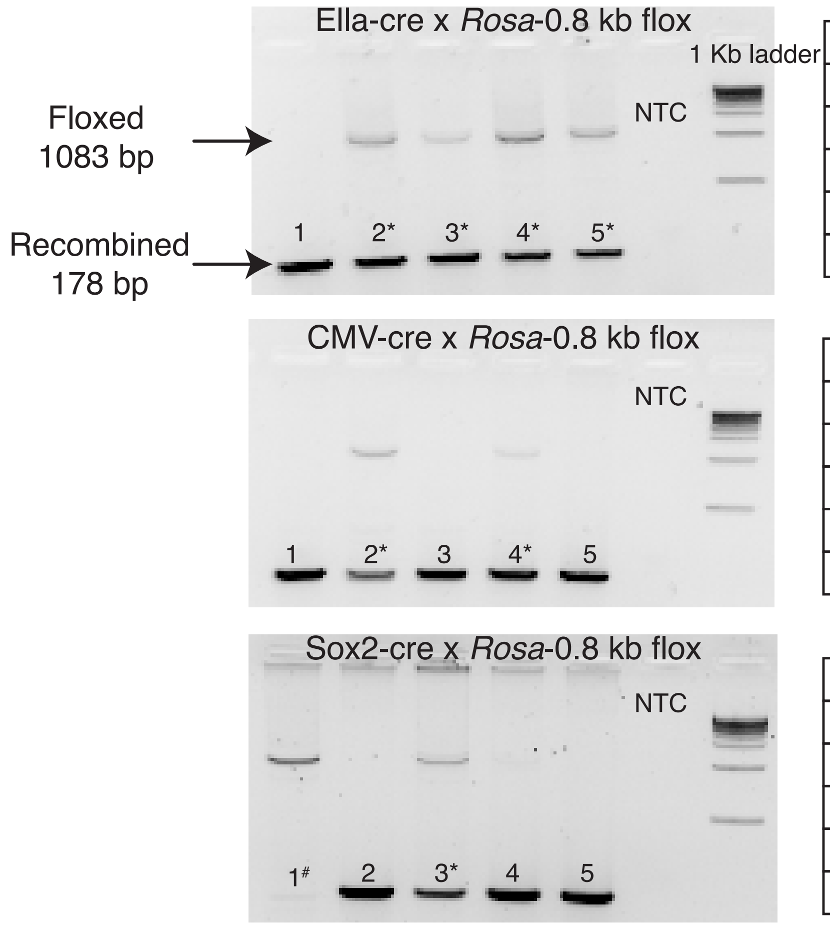


*Mosaic
#still floxed

**b**

Ella-cre x *Rosa*-0.8 kb flox

| | Recombined | Floxed | Rosa26 |
|---|---|---|---|
| Liver | + | + | + |
| Spleen | + | + | + |
| Kidney | + | + | + |
| Skin | + | + | + |
| Brain | + | + | + |

CMV-cre x *Rosa*-0.8 kb flox

| | Recombined | Floxed | Rosa26 |
|---|---|---|---|
| Liver | + | + | + |
| Spleen | + | + | + |
| Kidney | + | - | + |
| Skin | + | + | + |
| Brain | + | + | + |

Sox2-cre x *Rosa*-0.8 kb flox

| | Recombined | Floxed | Rosa26 |
|---|---|---|---|
| Liver | + | + | + |
| Spleen | + | + | + |
| Kidney | + | + | + |
| Skin | + | + | + |
| Brain | + | - | + |

### Supplemental Figure 2

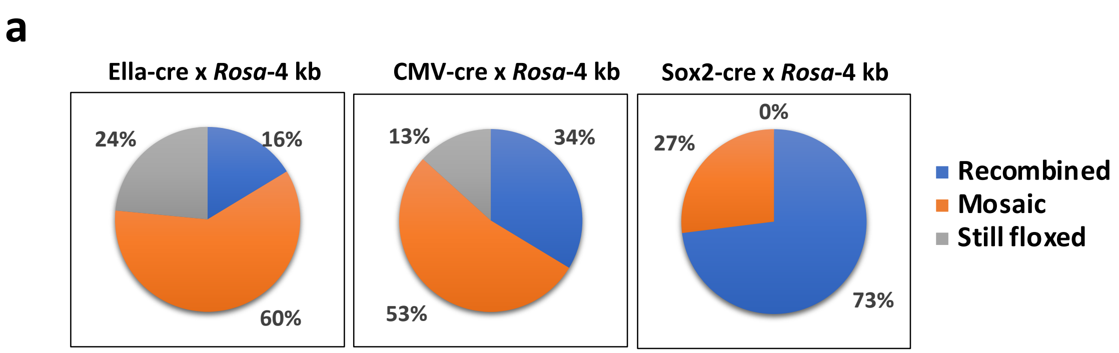


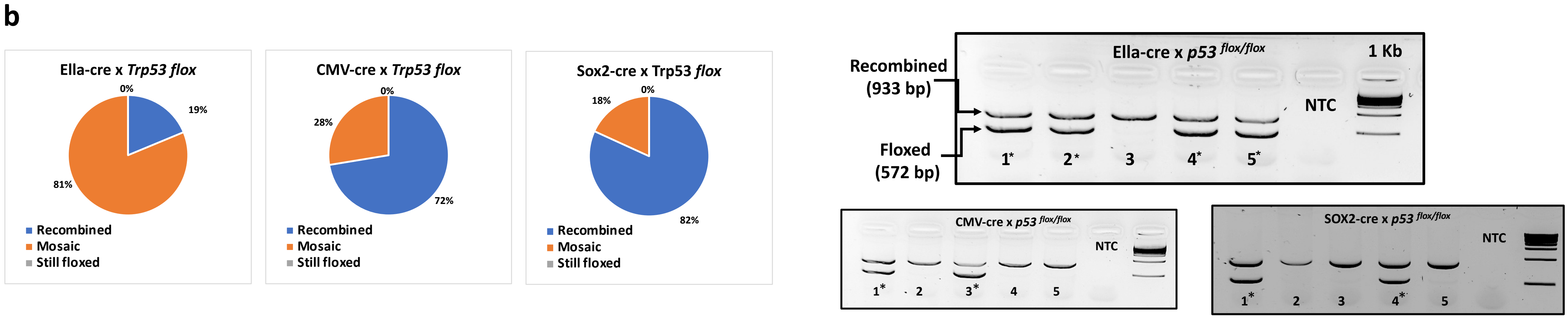

### Supplemental Figure 3

**a**

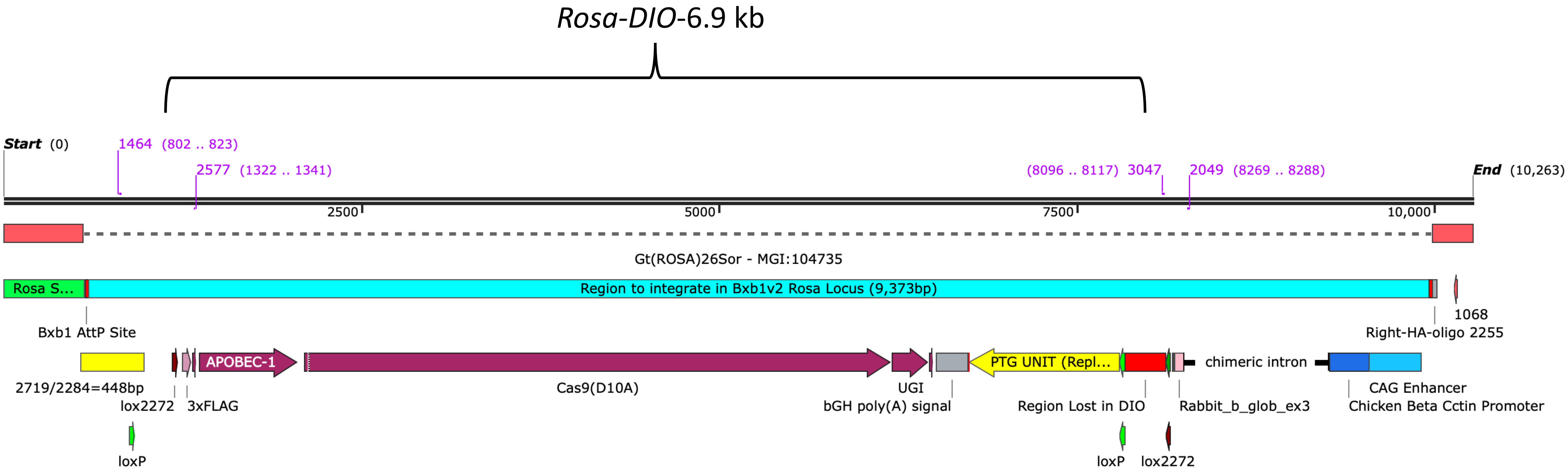


**b**

| Ella-cre X *R0SA-DIO* F1 offspring | |
|---|---|
| **Strain name** | **n** |
| Rosa-DIO-1.1 kb | 25 |
| Rosa-DIO-2.2 kb | 42 |
| Rosa-DIO-5.3 kb | 26 |
| Rosa-DIO-6.9 kb | 24 |
| Rosa-DIO-8.1 kb | 32 |

| CMV-cre X *R0SA-DIO* F1 offspring | |
|---|---|
| **Strain name** | **n** |
| Rosa-DIO-1.1 kb | 26 |
| Rosa-DIO-2.2 kb | 29 |
| Rosa-DIO-5.3 kb | 35 |
| Rosa-DIO-6.9 kb | 23 |
| Rosa-DIO-8.1 kb | 28 |

| Sox2-cre X *R0SA-DIO* F1 offspring | |
|---|---|
| **Strain name** | **n** |
| Rosa-DIO-1.1 kb | 43 |
| Rosa-DIO-2.2 kb | 14 |
| Rosa-DIO-5.3 kb | 26 |
| Rosa-DIO-6.9 kb | 24 |
| Rosa-DIO-8.1 kb | 55 |

#### Supplemental Figure 4

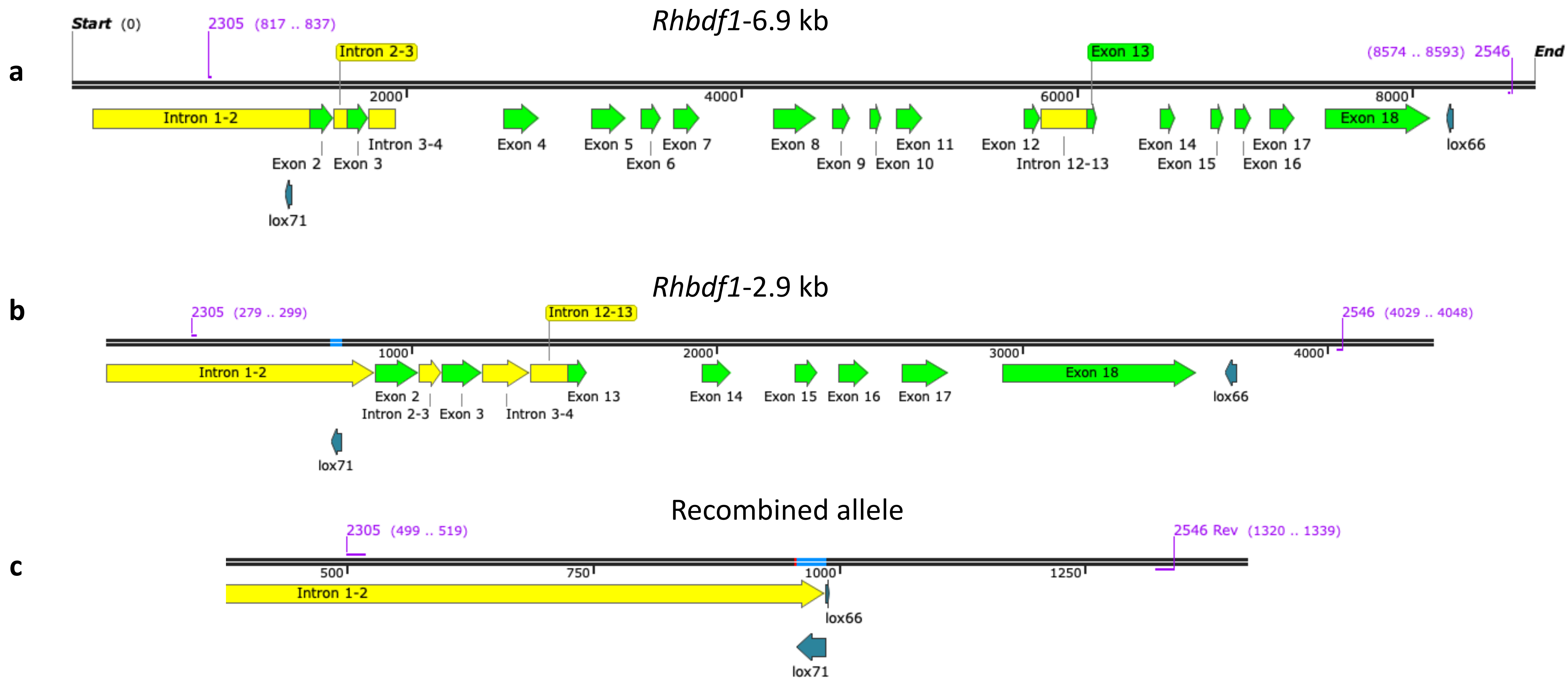

#### Supplemental Figure 5

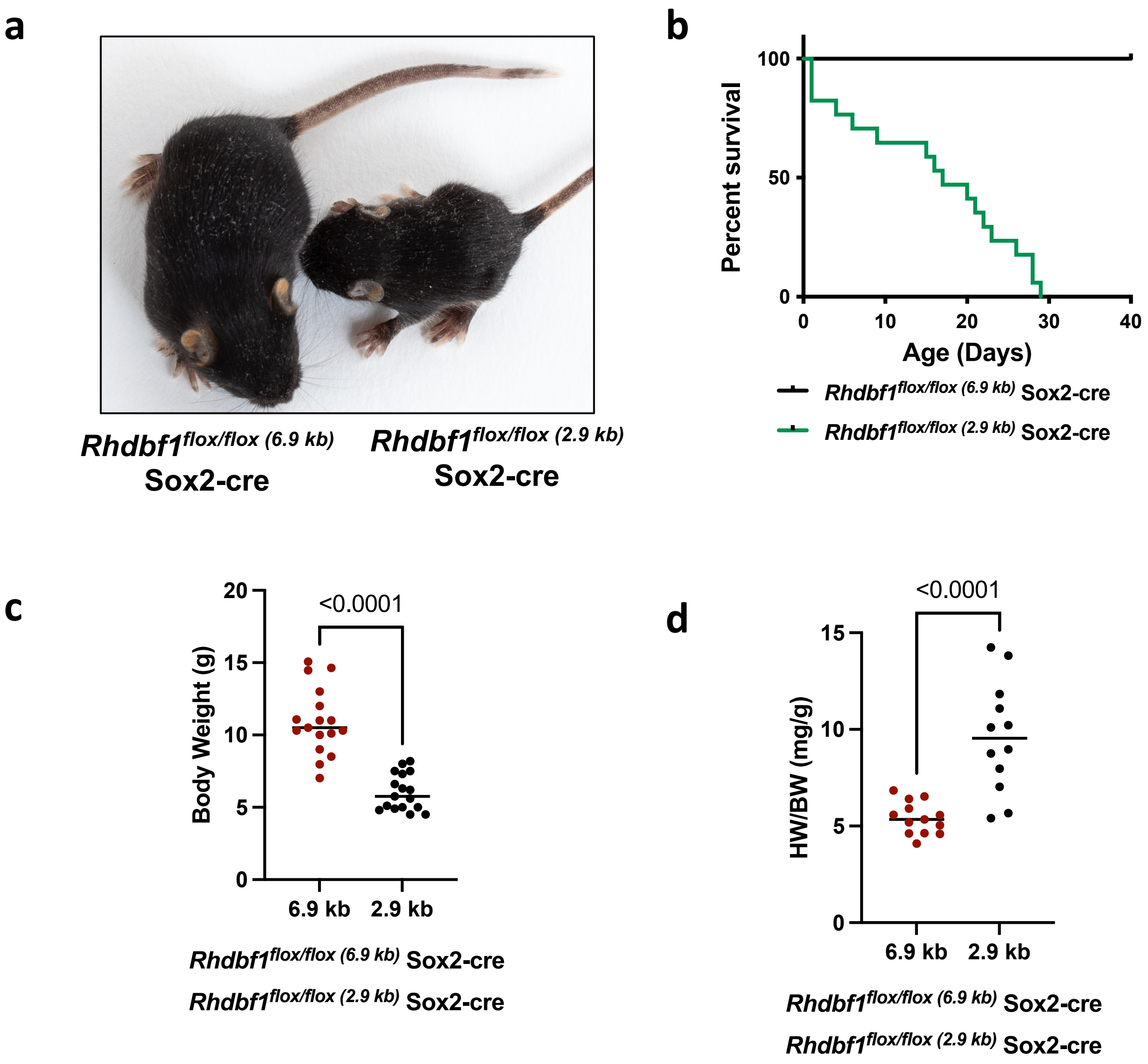

### Supplemental Figure 6

**a**

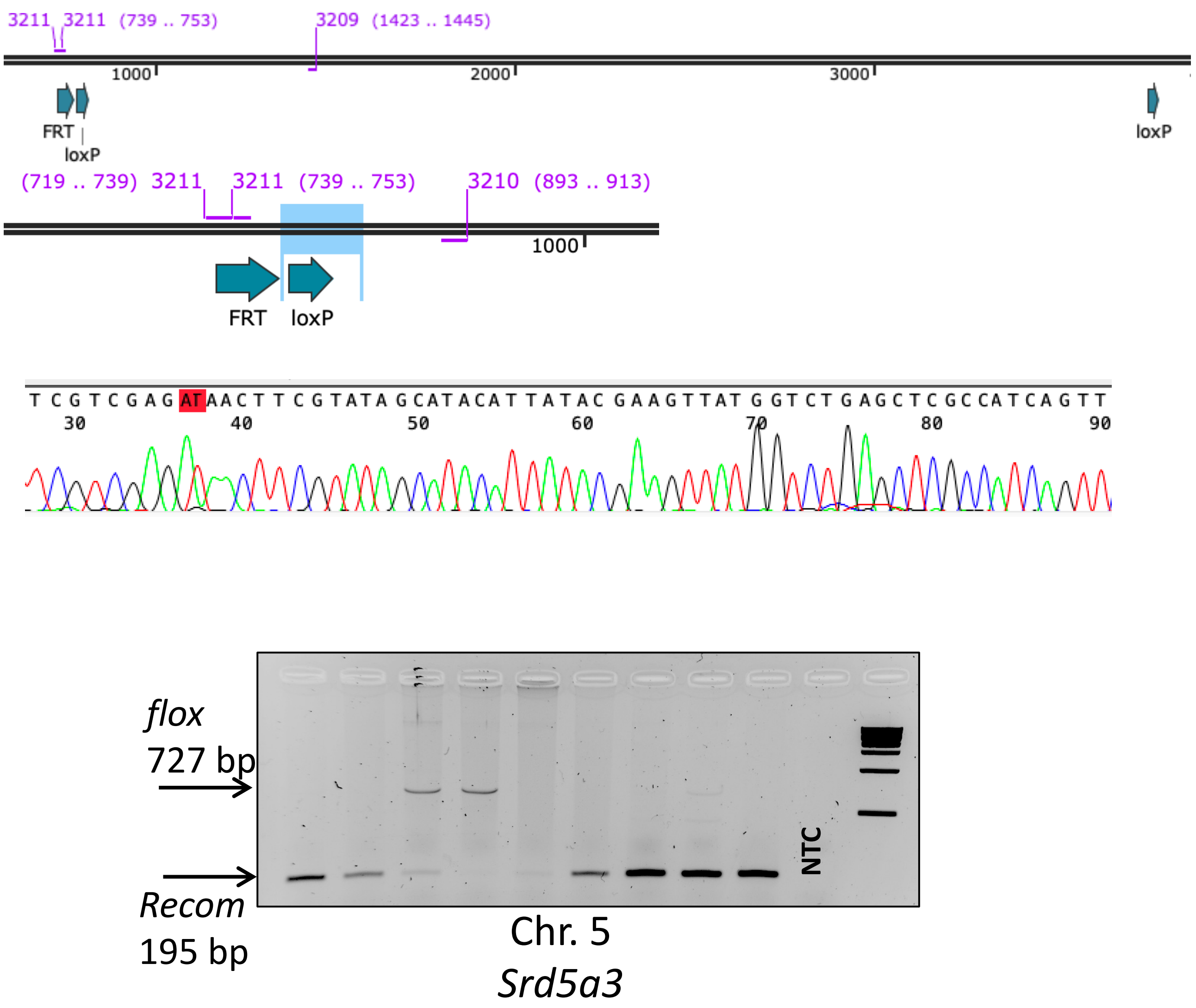


**b**

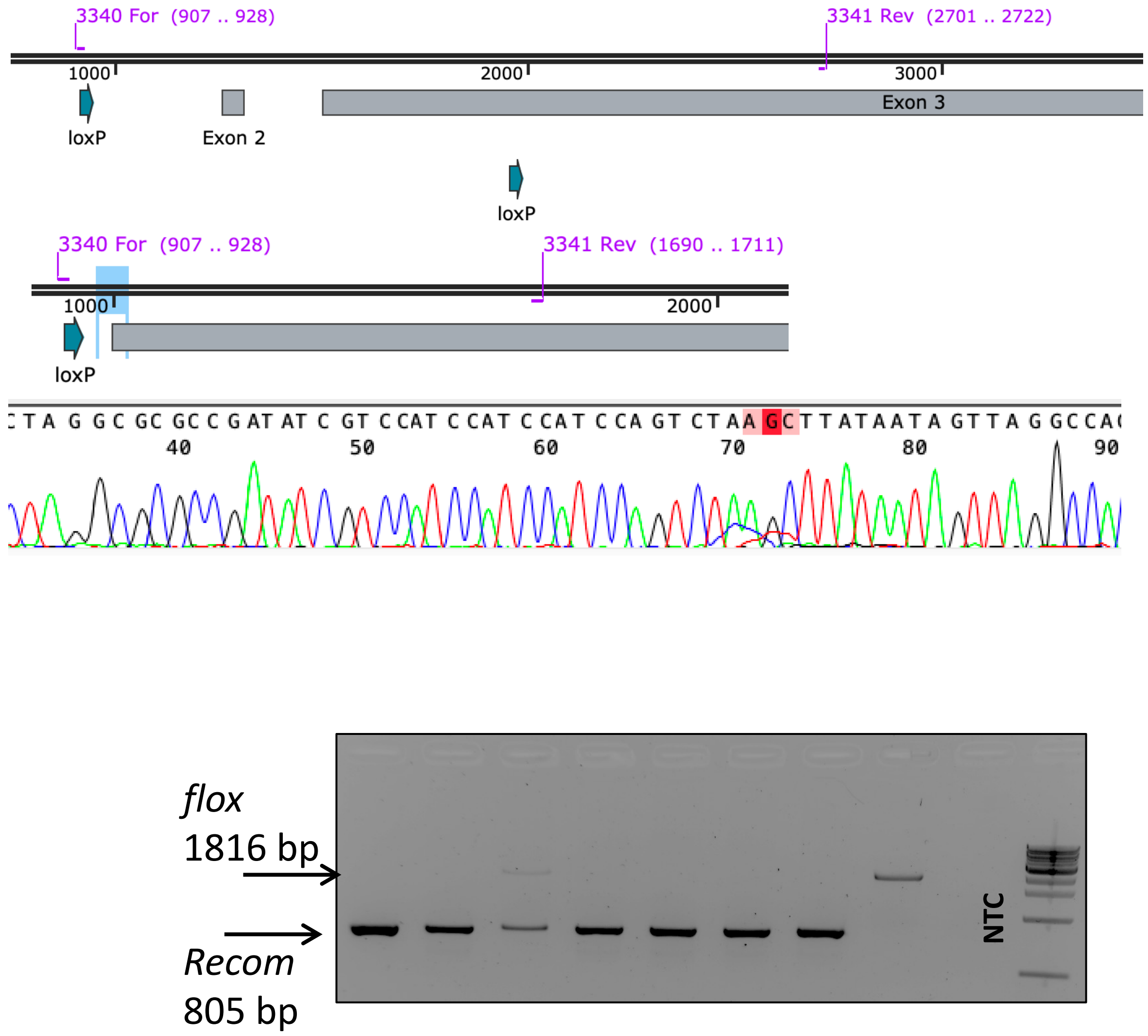

### Supplemental Figure 7

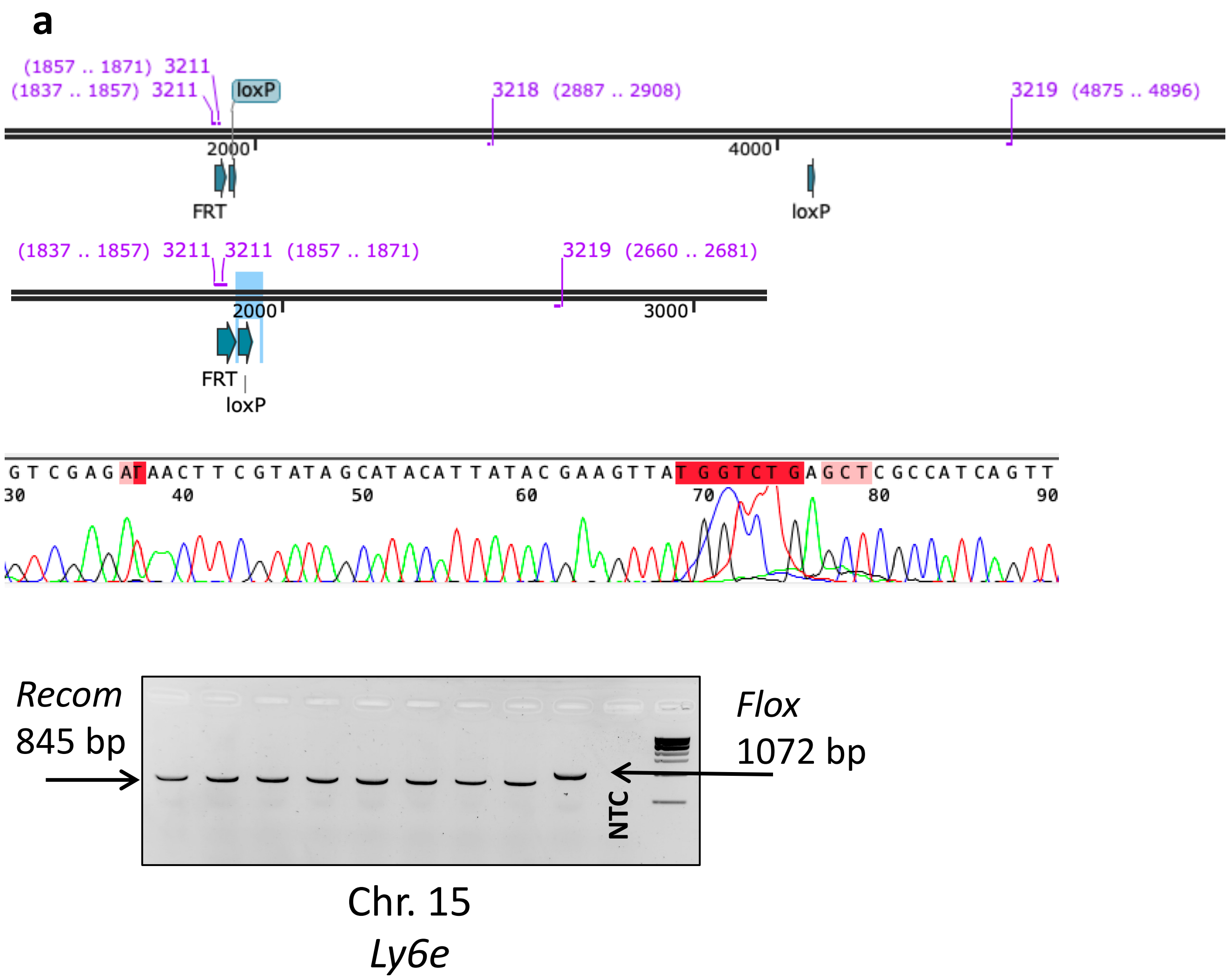


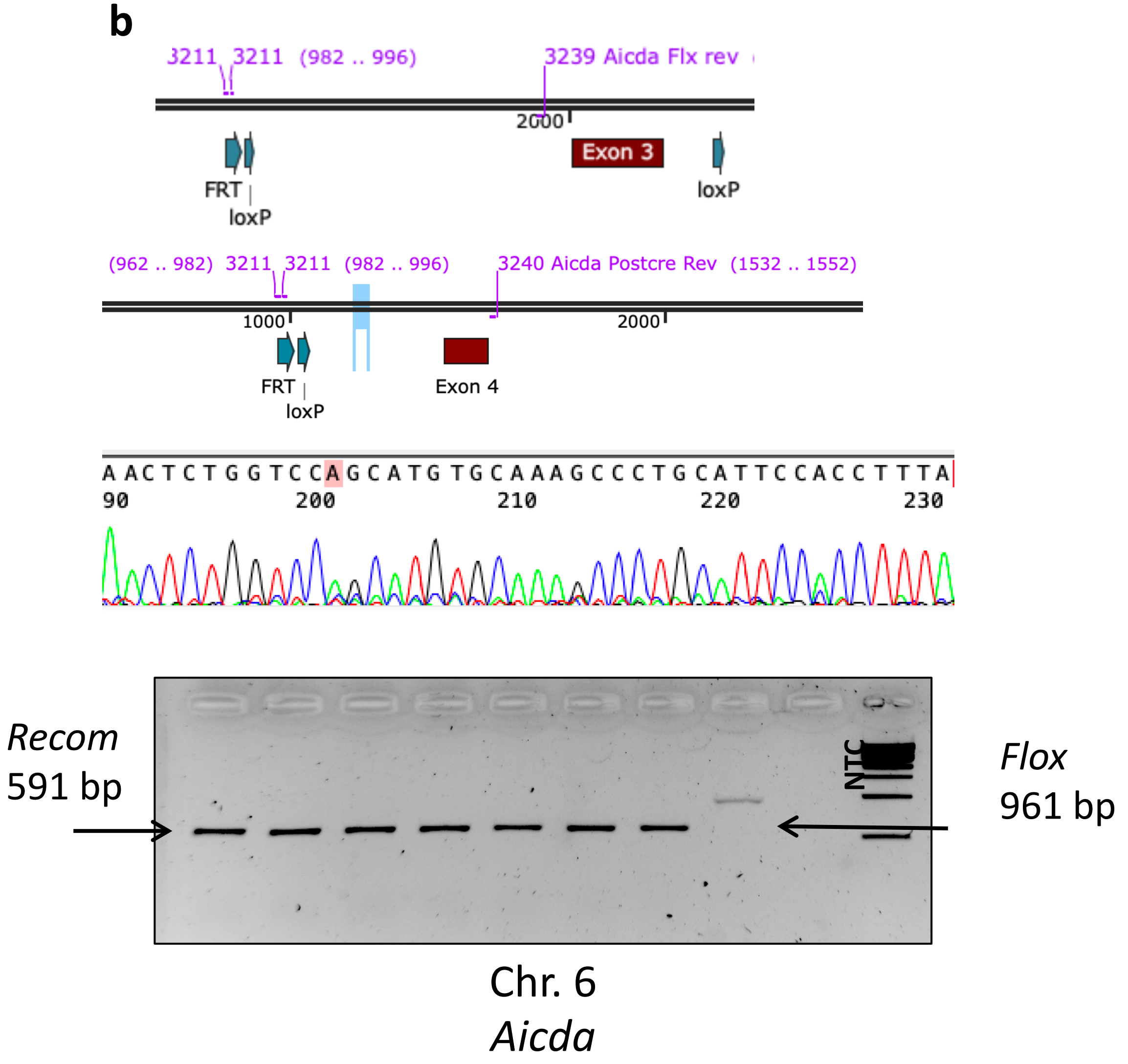

### Supplemental Figure 8

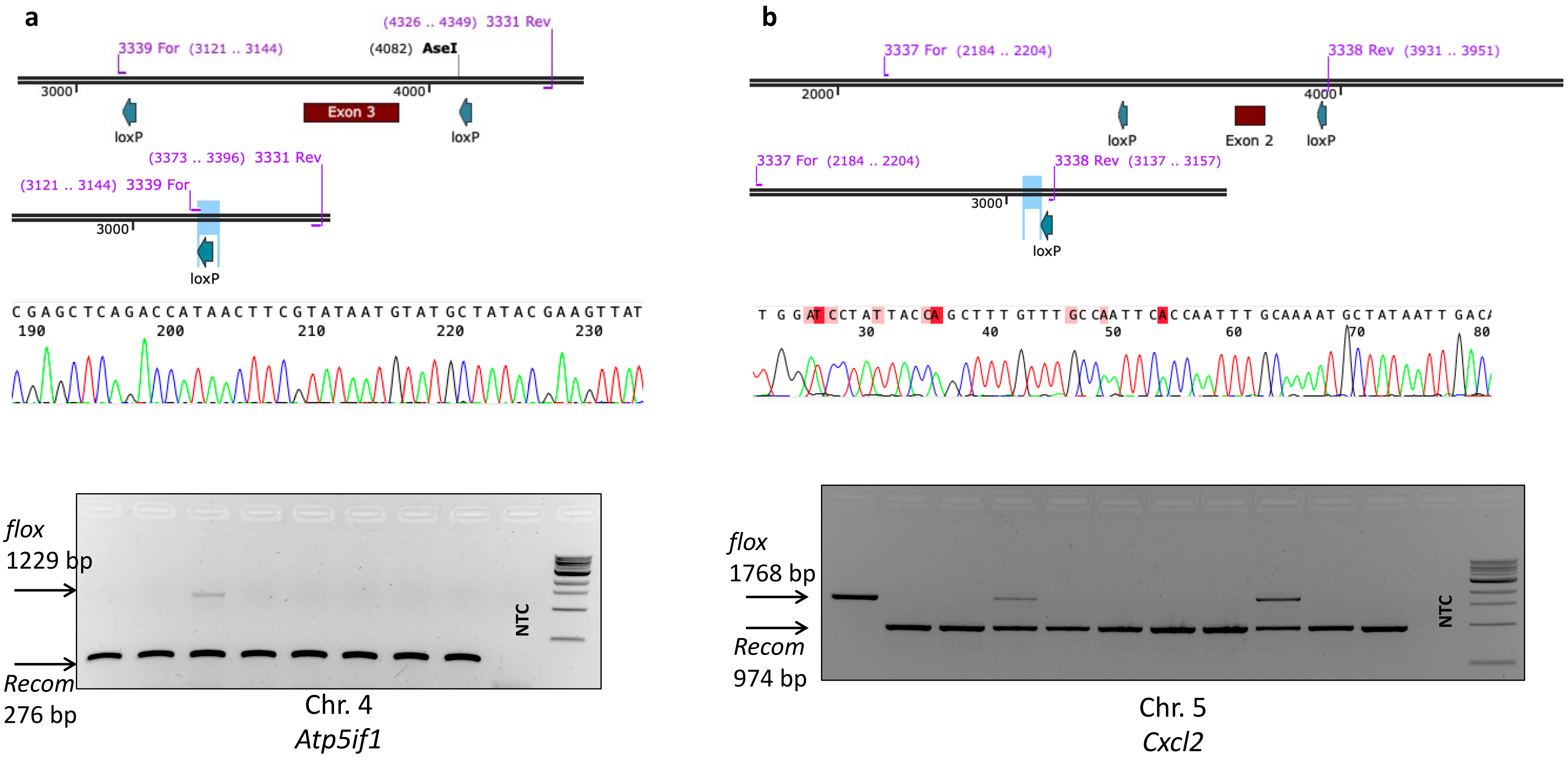

### Supplemental Figure 9

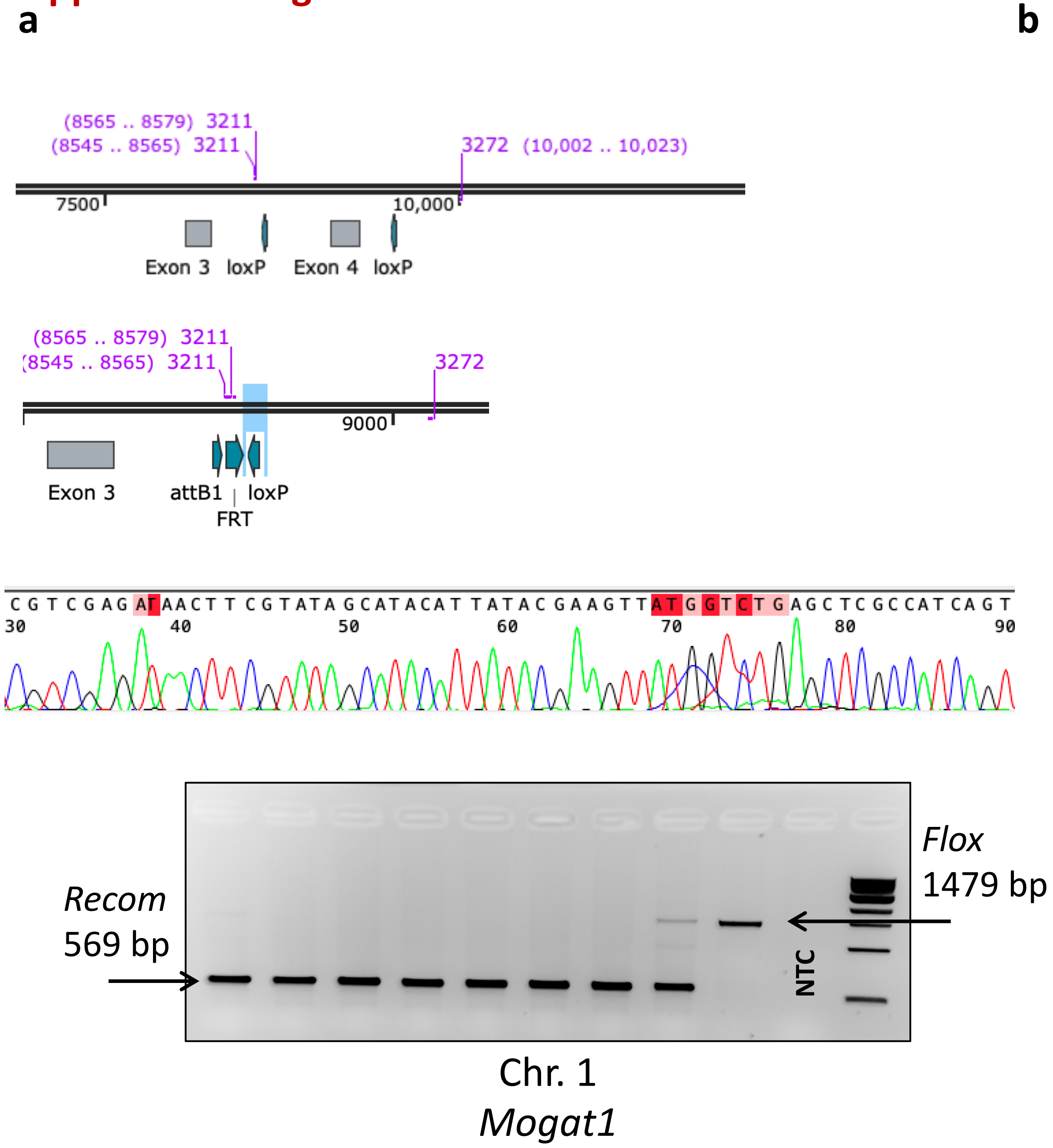


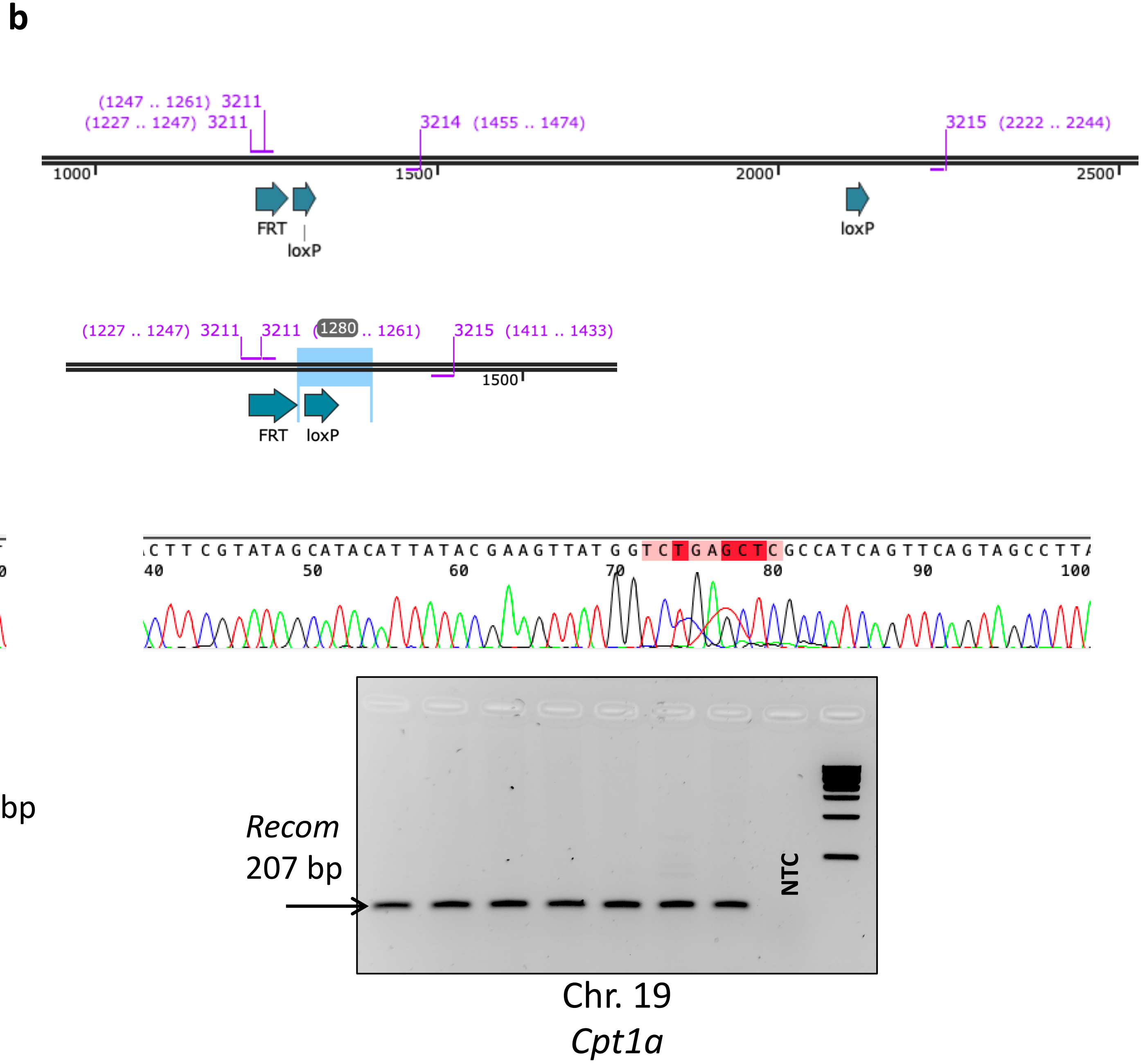

### Supplemental Figure 10

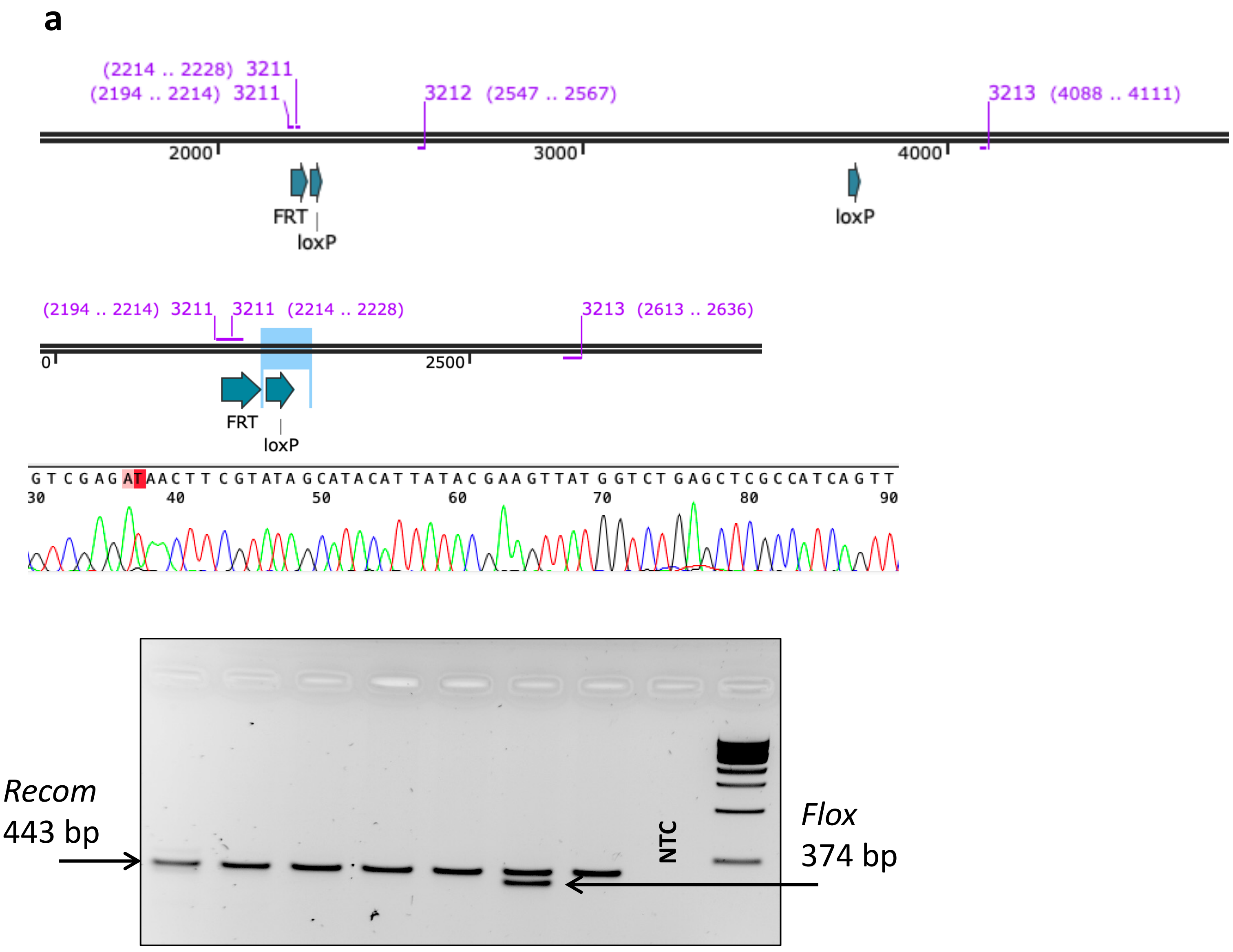


Chr. 8
*Nfatc3*

### Supplemental Figure 11

**a**

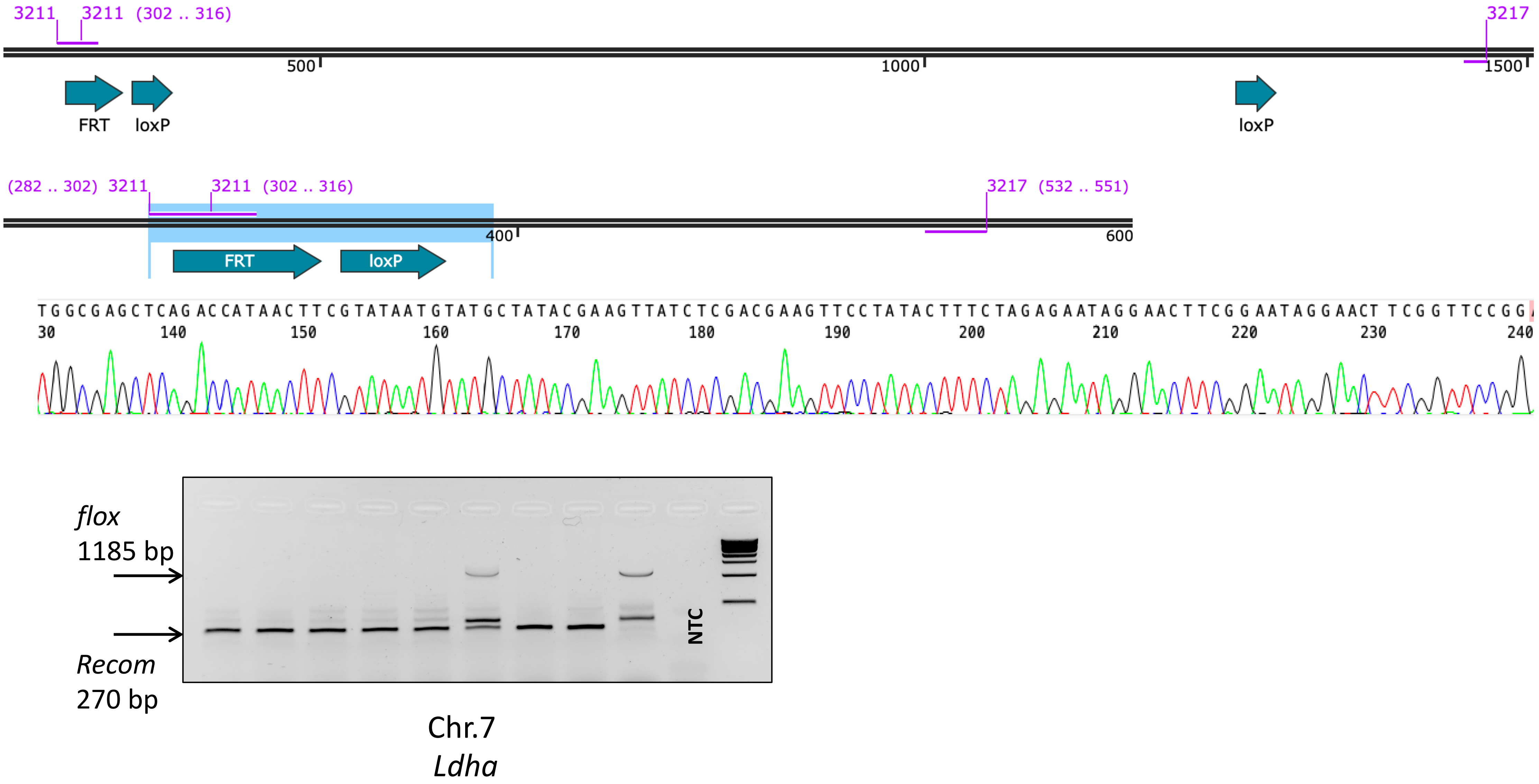

### Supplemental Figure 12

**a**

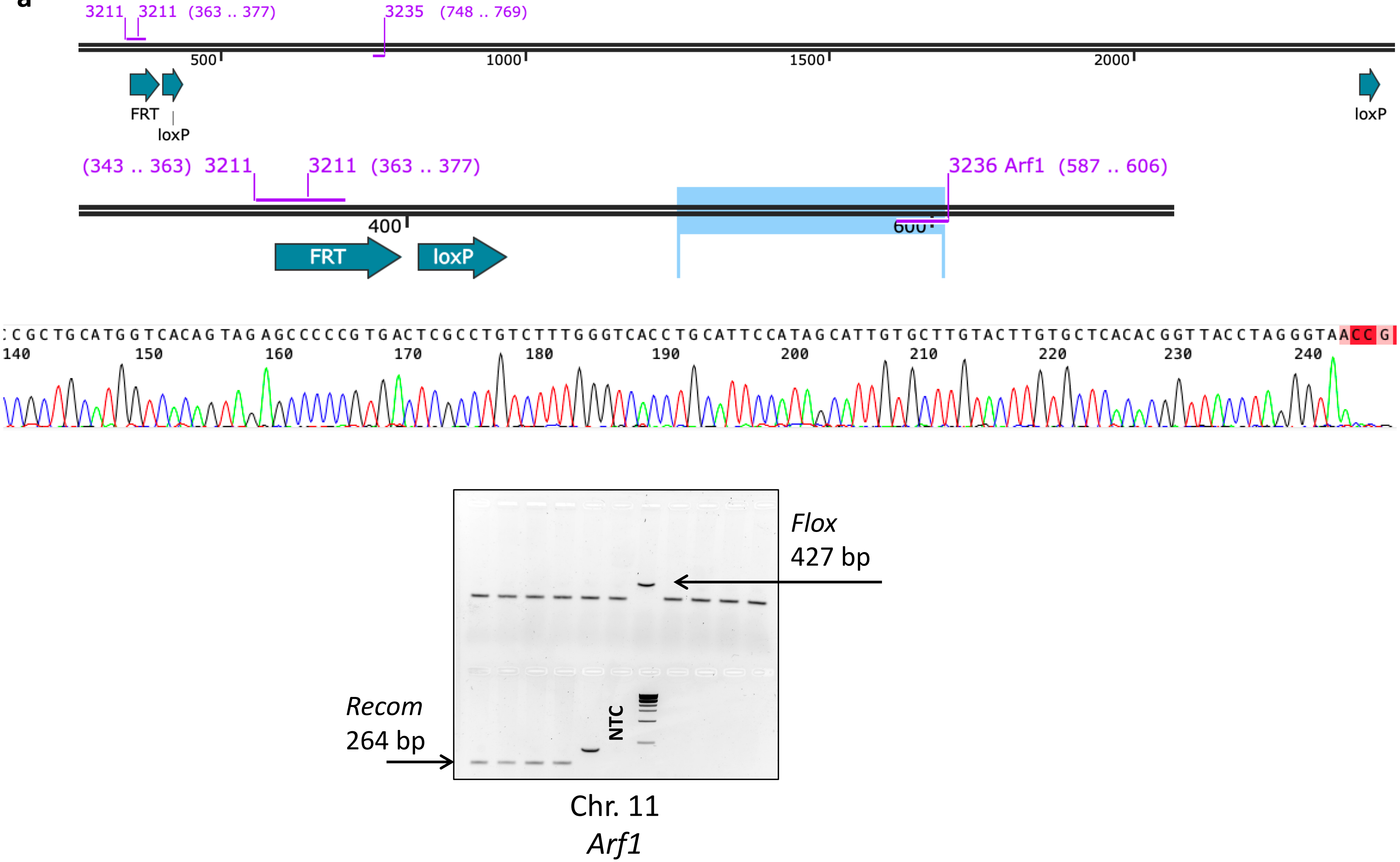

### Supplemental Figure 13

*2.7 kb Floxed Chr.11*

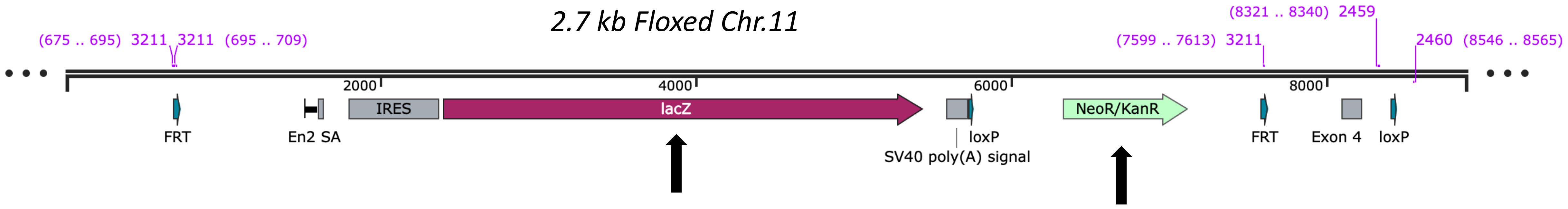


*Recombined*

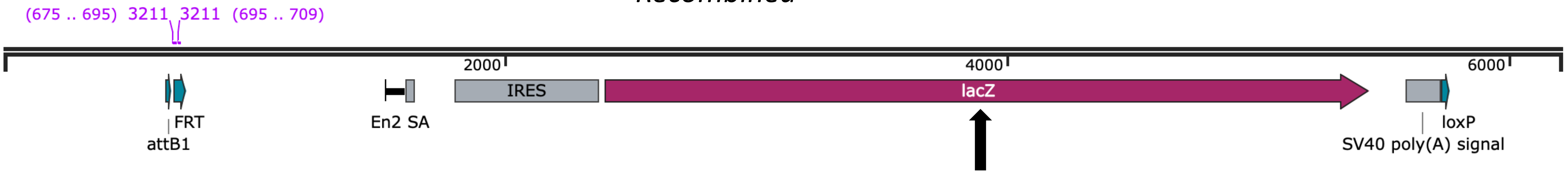

### Supplemental Figure 14

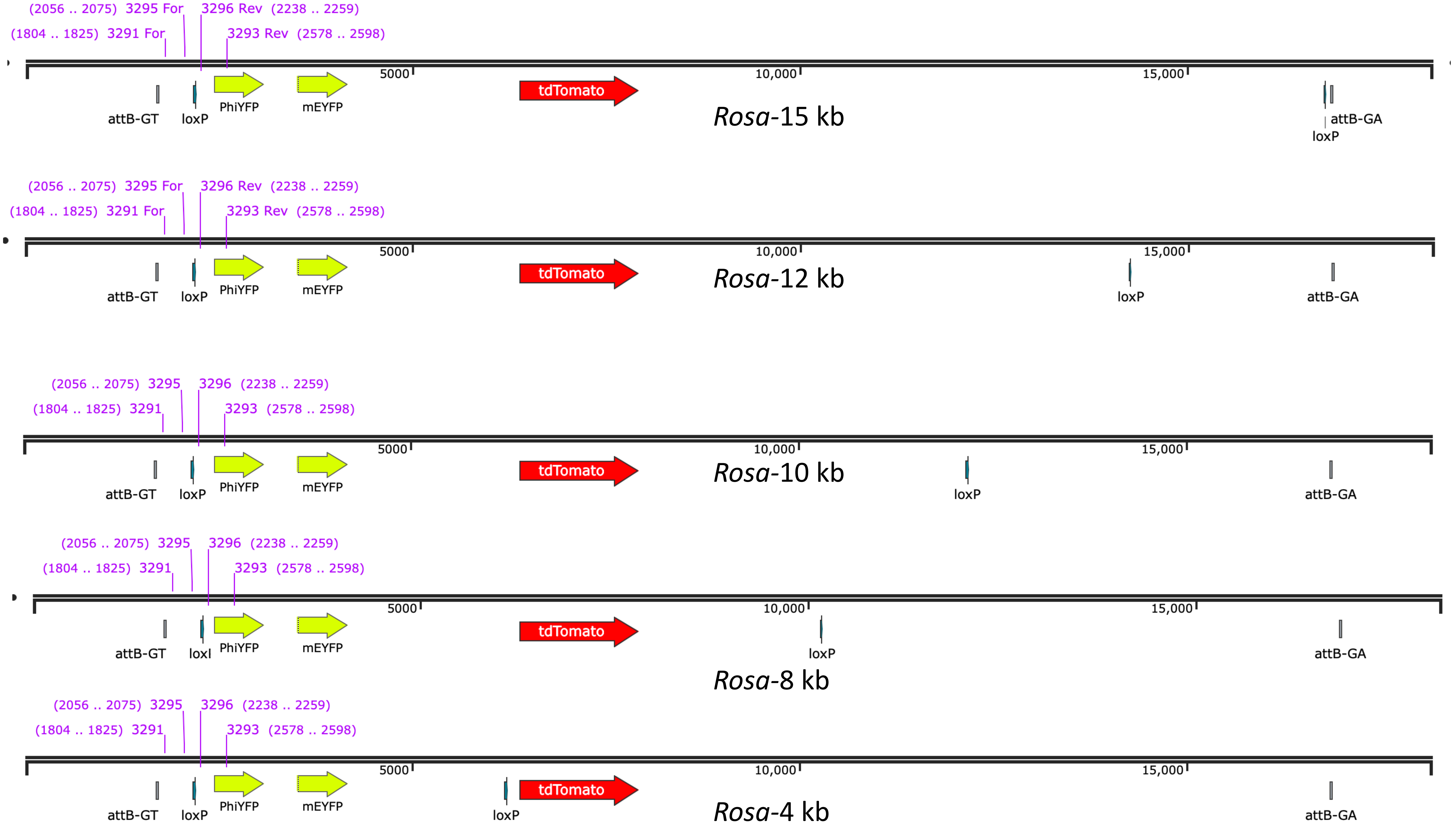
